# The nuclear pore complex connects energy sensing to transcriptional plasticity in longevity

**DOI:** 10.1101/2025.02.17.638704

**Authors:** Yifei Zhou, Fasih M Ahsan, Alexander A Soukas

## Abstract

As the only gateway governing nucleocytoplasmic transport, the nuclear pore complex (NPC) maintains fundamental cellular processes and deteriorates with age. However, the study of age-related roles of single NPC components remains challenging owing to the complexity of NPC composition. Here we demonstrate that the master energy sensor, AMPK, post-translationally regulates the abundance of the nucleoporin NPP-16/NUP50 in response to nutrient availability and energetic stress. In turn, NPP-16/NUP50 promotes transcriptomic activation of lipid catabolism to extend the lifespan of *Caenorhabditis elegans* independently of its role in nuclear transport. Rather, the intrinsically disordered region (IDR) of NPP-16/NUP50, through direct interaction with the transcriptional machinery, transactivates the promoters of catabolic genes. Remarkably, elevated NPP-16/NUP50 levels are sufficient to promote longevity and metabolic stress defenses. AMPK-NUP50 signaling is conserved to human, indicating that bridging energy sensing to metabolic adaptation is an ancient role of this signaling axis.

## Introduction

In response to reduced nutrient availability, animals rewire their metabolism for survival by promoting lipid catabolism and autophagy^1,2^. Nutritional stress such as that caused by dietary restriction (DR) impacts the activity of sentinel energy sensors, i.e., through activating AMP-activated protein kinase (AMPK) and inhibiting mTOR pathway. In turn, these sensors activate metabolic defense pathways, thereby promoting healthspan and longevity across species^3,4^.

Growing evidence suggests that low levels of nutritional stress encountered in early life can provide hormetic benefits across the entire lifespan and can even benefit progeny via transgenerational inheritance^5–7^. Multiple drugs and natural compounds tested or currently undergoing clinical trials to promote healthy aging also function through metabolic regulation, such as rapamycin, metformin, and NAD^+^ supplementation^8,9^. In some contexts, sensing nutrient deprivation is sufficient to modulate metabolism and lifespan even with abundant food availability^10^, highlighting the modulation of nutrient and energetic sensing pathways as an emerging strategy to intervene in aging. However, the full spectrum of mechanisms bridging energy sensing to modulation of aging remains incompletely characterized.

The nuclear pore complex (NPC) is a massive protein complex that is best known for its role in gating nucleocytosolic communication. The NPC consists of about 30 different proteins, known as nucleoporins, as a heteromultimeric assembly of more than 1000 protein subunits that mediate nuclear pore permeability, active transport, and on-site transcription^11–13^. In aging, both passive and active transport through the NPC are dysregulated, compromising nucleocytoplasmic trafficking, and causally contributing to age-related disorders^14–18^. For example, both in patients with Huntington’s disease and in murine models of the disease, nucleoporins are mislocated and aggregated with the disease-causing Htt protein in nuclei, compromising nucleocytoplasmic transport and neuronal fucntion^16^. Proper function of the NPC is critical to multiple pro-longevity paradigms that promote metabolic homeostasis, including reduced insulin/IGF-1 signaling (IIS)^14^, lysosomal lipase activation^19^, and metformin treatment^20^. Given the importance of the NPC in aging as well as the complex, multifaceted nature of the NPC’s composition, structure, and function, there is a critical need to determine granular mechanisms by which nucleoporins promote favorable effects in aging. Here we reasoned that a fuller understanding of how the NPC connects energy sensing to specific longevity effector mechanisms could illuminate multiple, heretofore unappreciated therapeutic inroads to promote healthy aging.

In this study, we demonstrate that the NPC subunit NPP-16/NUP50 bridges energy sensing and metabolic adaptation independently of its canonical role in nuclear permeability and transport. In response to energetic and nutrient stress, NPP-16/NUP50 is post-translationally activated by AMPK, and subsequently promotes the transcription of lipid catabolic genes. Overexpression of NPP-16/NUP50 is sufficient to induce lipid catabolism in both nematodes and mammalian cells. NPP-16/NUP50 overexpression robustly extends the lifespan in *C. elegans* by enhancing the transcriptional activity of the metabolic transcriptional regulators NHR-49/HNF4 and HLH-30/TFEB, driving lipid catabolism. Unlike scaffold nucleoporins, altered levels or activity of NPP-16/NUP50 do not affect nuclear transport and permeability; instead, increased NPP-16/NUP50 levels are necessary and sufficient to promote metabolic adaptation and longevity via interaction of its intrinsically disordered region (IDR) with the promoters of lipid catabolic genes. Our findings identify a heretofore unappreciated, conserved role of a specific nucleoporin in energy sensing and deployment of metabolic stress defenses against aging and further uncover a noncanonical role for nucleoporin IDRs in direct transcriptional regulation.

## Results

### NPP-16/NUP50 is required for the metabolic adaptation to nutrient stress

To identify nucleoporins required for modulation of metabolism in aging, we performed an RNAi screen in *C. elegans* targeting 20 nucleoporins to identify knockdowns which mitigate augmented expression of the fatty acid β-oxidation reporter *acs-2p::GFP* in response to starvation and phenformin treatment (a biguanide structurally related to metformin) that also prompts lifespan extension in *C. elegans*^21^. The *acs-2p::GFP* reporter is a widely used marker to indicate energetic stress in starvation, treatment with biguanides, and electron transport chain (ETC) inhibition, all of which extend the lifespan in *C. elegans*^22–26^. We found that most nucleoporin RNAi blocked the induction of *acs-2p::GFP* in response to 4 hours of starvation and treatment with 4.5 mM phenformin, including previously identified regulators of biguanide-mediated lifespan extension *npp-3/*NUP205 and *npp-21/*TPR^20^ (Figures 1A and S1). We also found that *npp-7*/NUP153, *npp-2*/NUP85, and *npp-16*/NUP50 RNAi blunt energetic stress-induced *acs-2p::GFP* expression (Figure S1C). Of interest, *npp-7*/NUP153 and *npp-16*/NUP50 are both nuclear basket components and interact with each other^27^, but we found that *npp-16*/NUP50 has no obvious role in NPC permeability and worm development (Figure S1D), which contrasts sharply with strict requirement for *npp-7*/NUP153^28–30^. These findings imply that *npp-16*/NUP50 may play a role in promoting metabolic adaptation to energetic and nutrient stress independently of potential regulatory roles in NPC permeability and nuclear transport.

**Figure 1.**
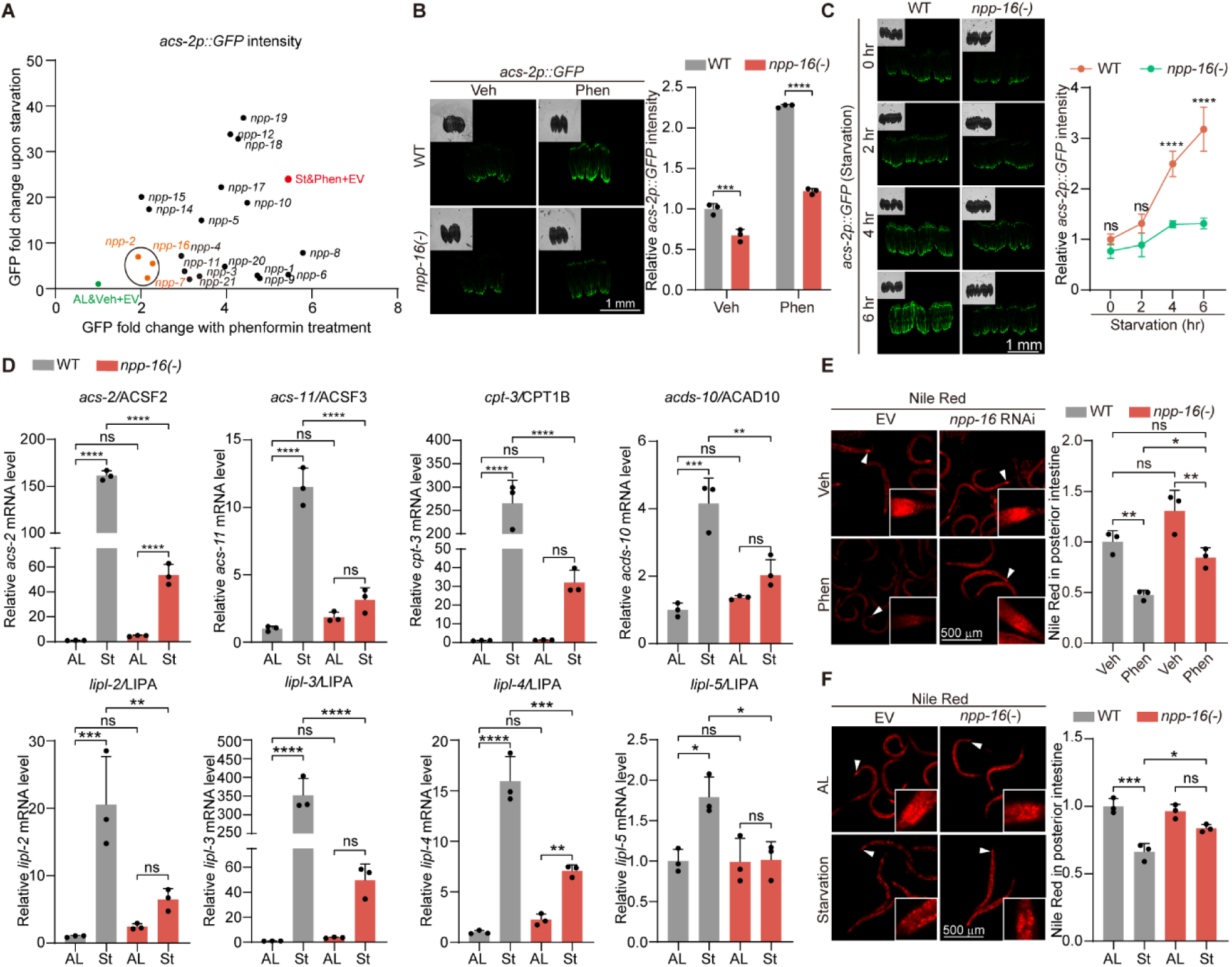
NPP-16/NUP50 is required for metabolic rewiring upon nutrient stress. **A.** An RNAi screen of 20 nucleoporins on *acs-2p::GFP* reporter expression upon starvation (St) and 4.5 mM phenformin treatment (Phen). GFP intensity is normalized to the empty vector (EV) RNAi group treated with vehicle (Veh) and *ad libitum* feeding (AL) as the fold change on the X and Y-axis, respectively. Starvation was performed for 4 hours at L4 before imaging. Phen and Veh treatment was performed from hatching to L4. The red font indicates the positive control (St&Phen+EV), the green font indicates the negative control (AL&Veh+EV), and the hits blunting *acs-2p::*GFP induction in both scenarios are highlighted by orange and circled. n=3 independent experiments. **B.** *acs-2p::GFP* reporter induction by 4.5 mM phenformin treatment is blocked by *npp-16(-)* loss of function. Worms were treated with Veh or Phen from hatching to day 1 adulthood (D1). Scale bar: 1 mm. n=3 independent experiments. **C.** The induction of *acs-2p::GFP* by starvation for the indicated duration is inhibited by *npp-16(-)* loss of function at D1. Scale bar: 1 mm. n=3 independent experiments. **D.** qRT-PCR analysis of lipid catabolic gene expression reveals that *npp-16(-)* loss of function rescues the induced mRNA level of lipid catabolic genes by starvation for 4 hours at L4. n=3 independent experiments. **E.** Neutral lipid storage indicated by fixative Nile red staining reveals that *npp-16(-)* loss of function inhibits lipid mobilization prompted by 4.5 mM phenformin in posterior intestinal cells at D1, indicated by white arrowheads and enlarged in white boxes. Scale bar: 500 μm. n=3 independent experiments. **F.** Neutral lipid storage indicated by fixative Nile red staining reveals that *npp-16(-)* loss of function abolishes the fat mass loss induced by starvation of 6 hours in the posterior intestinal cells at D1. Scale bar: 500 μm. n=3 independent experiments. Bars represent mean ± SD. Statistical significance was determined by two-way ANOVA relative to wild type (WT) worms (B and C) and the one-way ANOVA (D-F). Relative mRNA levels were normalized to *act-1* (D). ns: non-significant, \**p*<0.05, \*\**p*<0.01, \*\*\**p*<0.001, \*\*\*\**p*<0.0001.

We further investigated the function of *npp-16* in metabolic rewiring upon energetic and nutrient challenge. Upon food deprivation, activation of lipid catabolism and fatty acid β-oxidation enhances acetyl-CoA production to fuel the tricarboxylic acid cycle and ketogenesis, thereby providing alternative fuels for survival and fundamental cellular processes^31,32^. By quantitative reverse transcription PCR (qRT-PCR), we found that a group of catabolic genes, involved in lipolysis and lipid β-oxidation, are induced by starvation for 4 hours as reported^22,33^, including *acs-2* and *acs-11*, which are the orthologs of fatty acyl-CoA synthetase; *cpt-3*, an ortholog of carnitine palmitoyltransferase I; *acds-10*, which encodes an acyl-CoA dehydrogenase; and *lipl-2*, *lipl-3*, *lipl-4* and *lipl-5*, which encode lysosomal lipases in *C. elegans* (Figure 1D). Importantly, a null mutation of *npp-16* (*npp-16(-)*) ablates or significantly attenuates the transcriptional activation of these energetic and nutrient stress response genes (Figure 1D). Consequently, while wild type organismal fat mass is reduced upon starvation and phenformin treatment as a result of enhanced lipid catabolism^25,34^, *npp-16(-)* mutation inhibits fat mass reduction by short-term starvation and phenformin treatment concordant with its effects on suppressing lipid catabolic gene expression (Figures 1E and 1F). These data reveal that *npp-16*/NUP50 is necessary to activate lipid catabolism upon exposure to energetic and nutrient stresses.

### Activation of AMPK leads to phosphorylation and post-translational increase in NPP-16/NUP50 abundance

We next examined the expression pattern of NPP-16/NUP50 upon energetic stress. To visualize endogenous levels and localization of NPP-16, and to control its expression conditionally, we knocked in an in-frame, N-terminal GFP::AID::3xFLAG tag into the genomic *npp-16* locus using CRISPR/Cas9 technology^35,36^. By super-resolution confocal microscopy, we found the GFP-tagged endogenous NPP-16 is expressed ubiquitously in worms, and located on the nuclear envelope (NE) and in the nucleoplasm, consistent with mammalian NUP50 as previously reported^37,38^ (Figure S2A). Phenformin treatment and food deprivation significantly increase the fluorescence of endogenous NPP-16 on the NE and in the nucleoplasm of intestinal cells (Figures 2A and 2B). The protein level of NPP-16, as quantified by whole worm lysate immunoblotting, also increases significantly under these two scenarios (Figures 2C and 2D), whereas its RNA level is not elevated (Figure S2D). In contrast, CRISPR mCherry-tagged endogenous NPP-7/NUP153, which is responsible for anchoring NPP-16/NUP50 onto the NPC^27^ and also required for adaptive *acs-2p::GFP* induction (Figures 1A and S1), is unchanged by starvation (Figure S2B). mKate2::AID::3xFLAG-tagged endogenous NPP-11/NUP62, another nucleoporin at the central channel of the NPC, is also not affected by starvation (Figure S2C), suggesting that the specific up-regulation of NPP-16/NUP50 does not simply result from a global increase in NPC number or as a consequence of CRISPR editing nucleoporins to include fluorescent epitope tags. Concordantly, NPP-16/NUP50 protein is also increased when electron transport chain (ETC) activity and ATP production are disrupted by *nuo-6* (Complex I) and *cco-1* (Complex IV) knockdown (Figure S2E), without altering its mRNA expression (Figure S2F). These data indicate that NPP-16/NUP50 protein level is elevated by energetic and nutrient stresses, most likely by post-transcriptional mechanisms.

**Figure 2.**
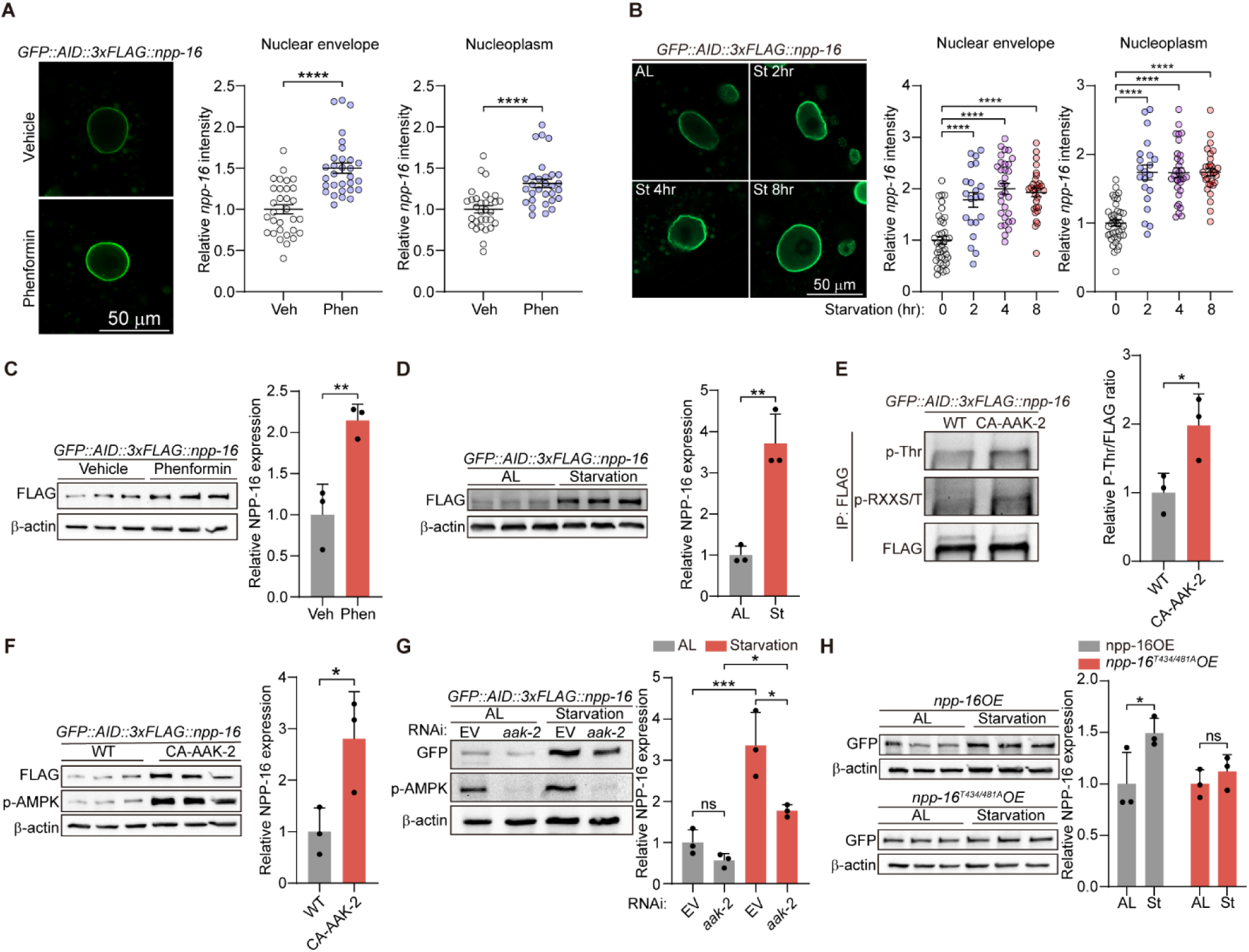
AMPK activity leads to increased phosphorylation and abundance of NUP50/NPP-16 protein. **A-B.** The fluorescence of endogenous *GFP::AID::3xFLAG::npp-16* is increased on the nuclear envelope (NE) and in the nucleoplasm of anterior intestinal cells when the worms are treated with 4.5 mM phenformin (Phen) and starvation (St) for the indicated times at L4. n=3 independent experiments containing at least 30 worms (A) and 21 worms (B) respectively. **C-D.** Endogenous protein levels of GFP::AID::3xFLAG::NPP-16 are induced by Phen treatment and starvation for 4 hours at L4. n=3 independent experiments. **E.** The level of phosphorylated threonines (p-Thr) and RxxS/T (RxxS/T-p) peptide motifs in immunoprecipitated endogenous NPP-16 is increased by overexpression of constitutively activated AAK-2 (CA-AAK-2) versus wild type animals (WT). The p-Thr signal is normalized to the signal of immunoprecipitated NPP-16. n=3 independent experiments. **F.** CA-AAK-2 increases the endogenous protein level of GFP::AID::3xFLAG::NPP-16 by immunoblotting at the L4 larval stage. n=3 independent experiments. **G.** Immunoblotting reveals that RNAi of *aak-2*/AMPKα abolishes the protein induction of NPP-16 by starvation for 4 hours at the L4 larval stage. n=3 independent experiments. **H.** Immunoblotting indicates that wild-type NPP-16 (*npp-16OE*) protein levels are significantly induced following 4-hours of starvation and are abolished by T434/481A mutation (*npp-16^T434/481A^OE*). n=3 independent experiments. Scale bar: 50 μm. Bars represent mean ± SEM (A-B) and mean ± SD (C-H). Statistical significance was calculated by unpaired *t*-test (A and C-F) and one-way ANOVA (B, G, and H). Relative protein levels were normalized to β-actin (C, D, and F-H). Empty vector (EV) serves as the negative control for RNAi experiments. ns: non-significant, \**p*<0.05, \*\**p*<0.01, \*\*\**p*<0.001, \*\*\*\**p*<0.0001.

AMP-activated protein kinase (AMPK) is an evolutionarily conserved, master hub of energy sensing and metabolic adaptation that coordinates cellular responses downstream of biguanide treatment, starvation, and ETC inhibition^1,39,40^. We thus suspected that AMPK might function upstream of NPP-16/NUP50 to post-translationally regulate its induction upon energetic stress. Compellingly, NPP-16/NUP50 harbors two conserved consensus sites for AMPK serine/threonine kinase phosphorylation ([M/I/L/V]XRXX[S/T]) at threonine 434 (T434) and threonine 481 (T481)^41^(Figure S2G), supporting the hypothesis that NPP-16/NUP50 may be a direct phosphorylation target of AMPK. As expected, in a worm strain overexpressing constitutively activated AAK-2 (CA-AAK-2)^42^, one of two *C. elegans* orthologues of AMPKα catalytic subunit, phosphorylation of both threonine (p-Thr) and serine/threonine within a RxxS/T (p-RXXS/T) basophilic kinase recognition motif are increased by CA-AAK-2 in immunoprecipitated NPP-16 vs. NPP-16 from wild-type worms (Figure 2E). Genetic activation of AMPK also increases the protein level of NPP-16 without altering its mRNA expression (Figures 2F and S2H). Moreover, *aak-2* knockdown by RNAi attenuates the induction of NPP-16 protein levels upon starvation (Figure 2G), indicating that nutrient stress increases NPP-16 in an AMPK-dependent manner. To further confirm the direct phosphorylation sites on NPP-16 that AMPK phosphorylates, we constructed integrated GFP-fusion strains overexpressing full-length NPP-16 (*npp-16OE*) and a T434/481A phospho-null NPP-16 variant which cannot be phosphorylated by AMPK on its conserved consensus sites (*npp-16^T4^*^34^*^/481A^OE*). While wild-type NPP-16 protein level is induced by starvation, NPP-16^T434/481A^ variant levels are unchanged (Figure 2H), demonstrating a firm requirement for consensus AMPK phosphorylation recognition sequences within NPP-16 for induction upon nutrient stress.

The AMPK consensus phosphorylation sites on NPP-16/NUP50 are conserved from *C. elegans* to human (Figure S2G), suggesting that human NUP50 expression might also be induced upon nutrient or growth factor deprivation. In keeping with this possibility, NUP50 is induced by serum starvation in HeLa cells, a nutrient stress known to activate AMPK, whereas NUP153 and NUP62 are unchanged (Figure S2I). This induction is suppressed by treating with compound C, an inhibitor of AMPK^43^ (Figure S2J), indicating that AMPK is the conserved upstream kinase that prompts the accumulation of NUP50 upon nutrient stress.

### NPP-16/NUP50 is required for the longevity paradigms associated with energetic stress

Interventions which deprive organisms of nutrients, such as caloric restriction, selective-nutrient restriction, or intermittent fasting, promote both healthspan and lifespan across a wide range of species^44–47^. Reduced activity of energy-sensing pathways such as reduced insulin/IGF-1 (IIS) and mTOR signaling also positively modulates lifespan and reduces age-related morbidity^3,4^.

We were therefore interested in whether the nutrient and energetic stress responsive activation of NPP-16/NUP50 also impacts longevity. We first examined a potential role for NPP-16/NUP50 in modulating biguanide-mediated lifespan extension, known to extend lifespan and health span in multiple organisms by inhibiting electron transport chain (ETC) activity and activating AMPK^39,48^. Consistent with the essential role of NPP-16 in lipid catabolism (Figure 1), *npp-16* RNAi suppressed the lifespan extension by phenformin treatment without altering the lifespan of wild-type worms treated with vehicle control (Figure 3A).

**Figure 3.**
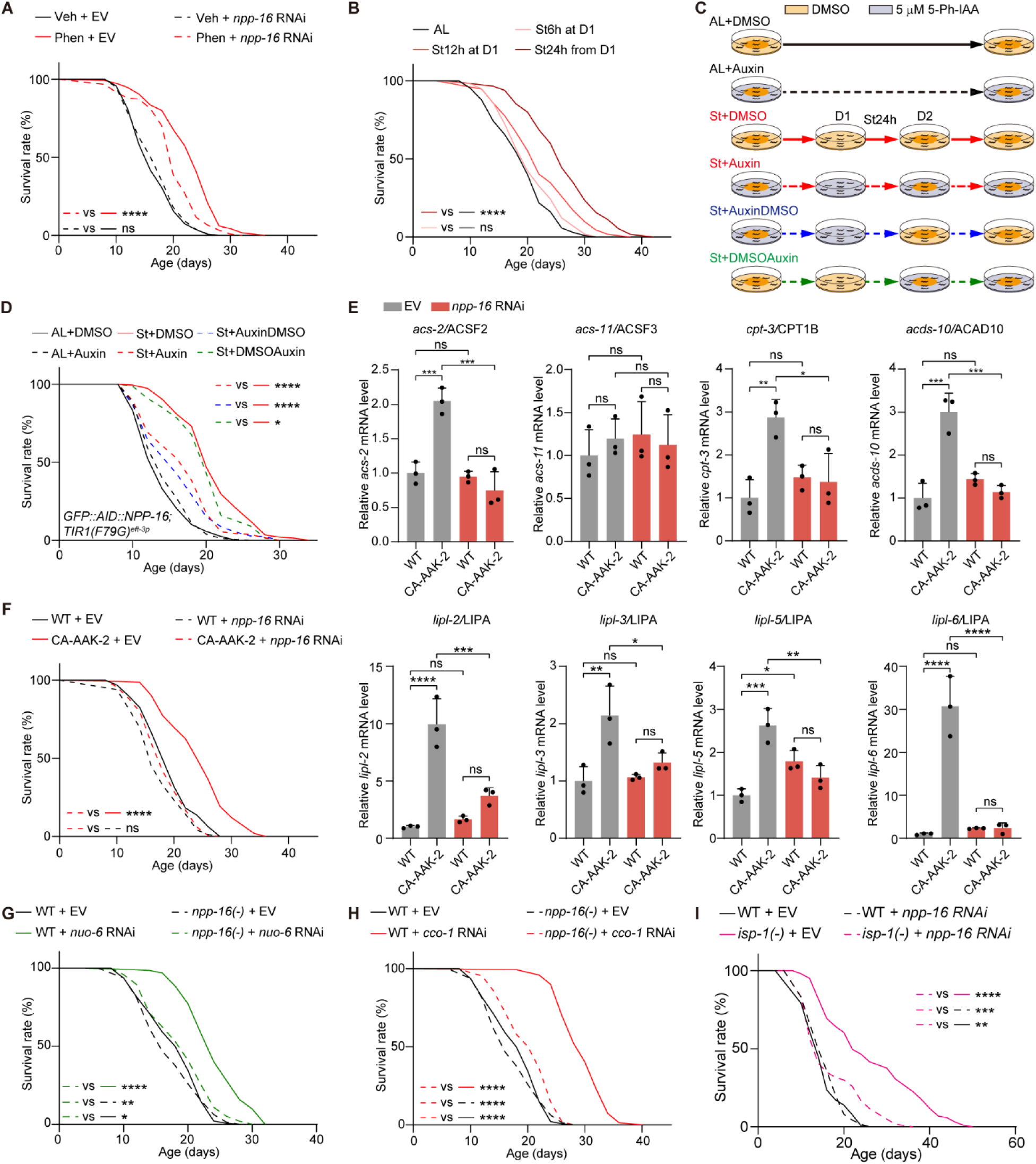
NPP-16/NUP50 activity is required for lifespan extension and metabolic rewiring across multiple pro-longevity paradigms. **A.** Knockdown of *npp-16* by RNAi inhibits the lifespan extension normally seen with phenformin treatment (Phen). Vehicle treatment (Veh) serves as the negative control for Phen. **B.** Lifespan analyses highlighting proof of principle lifespan extension from hormetic starvation (St) for indicated times at day 1 adulthood in wild-type nematodes. **C.** A schematic showing the design of the hormetic starvation for 24 hours (St24h) experiment with conditional knockdown of NPP-16 by the AID-TIR1(F79G) system. **D.** Conditional knockdown of NPP-16 by the AID-TIR1(F79G) system prior to refeeding suppresses the longevity phenotype seen following hormetic starvation, whereas conditional knockdown after starvation does not affect lifespan extension. **E.** qRT-PCR analyses reveal *npp-16* RNAi ablates the transcriptional up-regulation of lipid catabolic genes in CA-AAK-2 worms. n=3 independent experiments. **F.** *npp-16* is required for lifespan extension in CA-AAK-2 worms. **G-I.** *npp-1*6 is required for longevity seen following inhibition of electron transport chain (ETC) activity. The ETC is suppressed by either *nuo-6* RNAi, *cco-1* RNAi, or *isp-1(-)* mutation, which are components of ETC complexes I, IV, and III, respectively. Bars represented mean ± SD. Statistical significance was determined by one-way ANOVA (E). Empty vector (EV) serves as the negative control for RNAi experiments. Relative mRNA levels were normalized to *act-1* (E). ns: non-significant, \**p*<0.05, \*\**p*<0.01, \*\*\**p*<0.001, \*\*\*\**p*<0.0001. See also Table S1 for independent biological replicates and summary lifespan statistics.

Multiple distinct regimens of dietary restriction (DR) extend lifespan across species^3,4,49^. For example, hormetic starvation stress in early life is sufficient to promote longevity and delay age-related disorders^5,7,50,51^. As previously reported, worms subjected to hormetic starvation (St) for 12 and 24 hours at day 1 of adulthood and then refed *ad libitum* (AL) have significantly longer lifespans (Figure 3B). To determine whether NPP-16 functions in a critical capacity in the longevity that follows hormetic starvation stress, we conditionally knocked down NPP-16 by auxin-inducible degradation (AID). We degraded an endogenously CRISPR-tagged GFP::AID::3xFLAG::NPP-16 by exposing worms to 5 μM of the modified auxin 5-Ph-IAA at specified time points, activating the TIR1(F79G) mediated ubiquitin-proteasome system^35^. 5-Ph-IAA treatment from hatching to L4 leads to the degradation of ∼80% of endogenous NPP-16 (Figure S3A), and short-term treatment for 4 hours is also sufficient to deplete NPP-16 (Figure S3B). We found that *npp-16* knockdown across the lifespan prevents the pro-longevity phenotype of hormetic starvation (Figures 3C-3D). NPP-16 conditional knockdown before refeeding is also sufficient to inhibit lifespan extension, whereas knockdown starting from refeeding does not affect lifespan extension (Figures 3C-3D). In aggregate, these results indicate that the critical window for NPP-16 function in promoting hormetic starvation-prompted longevity is initial energy sensing during starvation and the immediate ensuing adaptive response, but not for long-term maintenance of the pro-longevity effect. We find that NPP-16 activity specifically regulates lifespan extension of AMPK-dependent DR regimens, as *npp-16* RNAi does not affect the longevity phenotype of *eat-2(-)* mutants (Figure S3C), an AMPK-independent DR model that compromises pharyngeal pumping^49^. Moreover, NPP-16 is not required for mutation mediated lifespan extension (Figure S3D), which is in contrast to reports for other nucleoporins such as NPP-7/NUP153 known to be important for lifespan extension in *daf-2(-)* mutants^14^, implying that nucleoporins can play distinct roles in different longevity paradigms. Genetic activation of AMPK increases starvation-induced lipid catabolic gene expression, an effect that is mitigated by *npp-16* knockdown by RNAi (Figures 3E and S3E).

Consistent with it playing an important role downstream of AMPK-mediated metabolic adaptation, *npp-16* is also necessary for the extended lifespan of transgenic worms bearing constitutively activated AAK-2 (Figure 3F). In aggregate, these data suggest that NPP-16/NUP50 is required for transcriptional reprogramming of lipid catabolism and subsequent lifespan extension in AMPK-dependent longevity paradigms.

The activity of the mitochondrial electron transport chain (ETC) dynamically regulates energetic status to meet the metabolic demands of cells and organisms^52^. Inhibition of ETC activity prompts metabolic adaption through activation of fatty acid β-oxidation^24^, which can be visualized by activation of the fatty acid β-oxidation reporter *acs-2p::GFP.* We found that *acs-2p::GFP* activation as well as induction of endogenous mRNAs for *acs-2* and *cpt-3* following knockdown of ETC components *nuo-6* and *cco-1* by RNAi are completely lost with *npp-16(-)* mutation (Figures S3F and S3G). However, *npp-16* is not required for the induction of the mitochondrial unfolded protein response (UPR^mt^) stress chaperone *hsp-6* by ETC inhibition (Figure S3G), suggesting that the action of NPP-16 is specific to metabolic adaptation.

Perturbation of ETC subunits also extends the lifespan in *C. elegans*^53^. Consistent with the requirement of *aak-2* for ETC inhibition mediated lifespan extension^54^, we found that *npp-16(-)* mutation suppresses the longevity phenotype of a broad-spectrum of genetic ETC inhibitions: *nuo-6* RNAi (complex I), *cco-1* RNAi (complex IV), and *isp-1(-)* mutation (complex III) (Figures 3G-3I). These results highlight a critical role for *npp-16*/NUP50 in the modulation of metabolic adaptation and longevity in response to energetic stress.

### Promotion of NUP50 expression is sufficient to extend lifespan by activating lipid catabolism

We next investigated whether overexpression of NPP-16/NUP50 is sufficient to promote longevity by altering lipid catabolic pathways, leveraging our *npp-16OE* worms, which have an approximate 4.7-fold increase in *npp-16* mRNA (Figure S4A). This level of *npp-16* overexpression alone is sufficient to upregulate expression of the *acs-2p::GFP* reporter and the mRNA levels of other lipid catabolic genes that respond to starvation and AMPK activation, phenocopying the transcriptional metabolic adaptation to energetic stress (Figures S4B, 4A, 1D and 3E). Indicating that this mechanism is conserved and ancient, we also found that human NUP50 overexpression (NUP50OE) in HeLa cells is sufficient to activate the expression of ACSF2, an acyl-CoA synthetase orthologous to *acs-2* in *C. elegans*, and significantly enhanced the induction of ACSF2, CPT1B, and LIPA upon serum deprivation relative to the cells transfected with EGFP empty vector (Figure 4B). As a result of activated lipid catabolism gene expression, *npp-16*OE animals also showed a significantly decreased fat mass compared to WT worms (Figure S4C), suggesting that driving *npp-16* overexpression can enhance functional lipid catabolism. Strikingly, *npp-16OE* extends lifespan by about 80% versus WT worms (Figure 4C), showing that NPP-16 is both necessary and sufficient to direct pro-longevity outcomes in *C. elegans*. As many energetic stress paradigms require lipolysis and lipid β-oxidation in worms to extend lifespan^26,55–57^, we then investigated whether *npp-16OE* requires lipid catabolism for lifespan extension. Indeed, knockdown of *acs-2* by RNAi abolishes the lifespan extension in *npp-16OE* worms (Figure 4D) and prevents fat mass loss (Figure S4D). RNAi knockdown of *cpt-3*, a rate-limiting enzyme for lipid β-oxidation^58^, also prevents the longevity phenotype of *npp-16OE* worms (Figure 4E), and knockdown of the lysosomal lipase *lipl-3* partially suppresses the longevity phenotype (Figure S4E), whereas *acs-11* RNAi has no obvious effect (Figure S4F).

**Figure 4.**
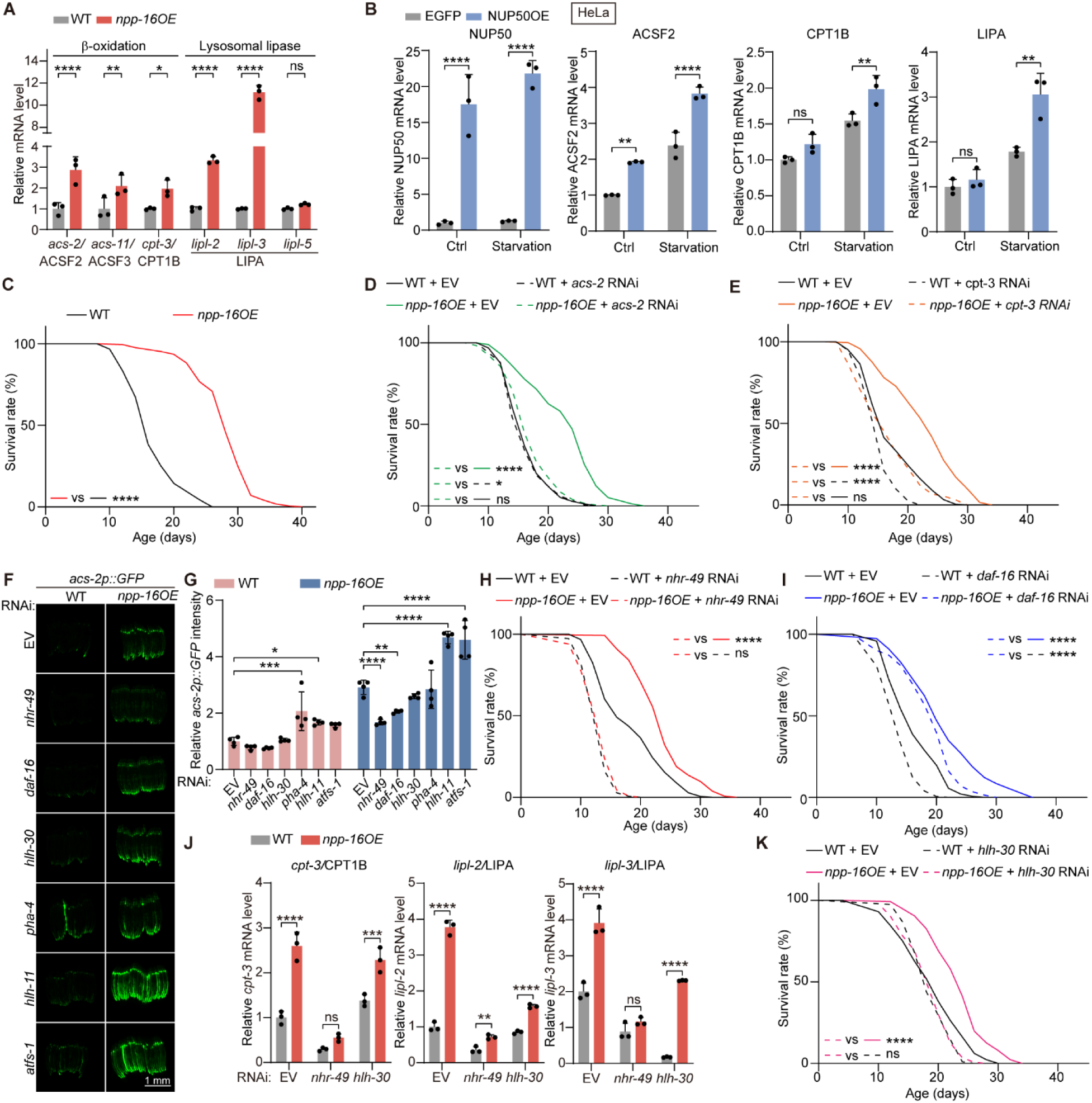
NPP-16/NUP50 overexpression promotes longevity through activating lipid catabolism. **A.** *npp-16OE* increases the abundance of mRNAs encoding lipid catabolic genes induced by either starvation or CA-AAK-2 by qRT-PCR. n=3 independent experiments. **B.** qRT-PCR analyses reveal that NUP50OE induces the abundance of mRNAs encoding lipid catabolic genes in HeLa cells in a complete medium (Ctrl) and serum-deprivation medium for 24 hours (Starvation). n=3 independent experiments. **C.** *npp-16OE* is sufficient to extend the lifespan of *C. elegans*. **D-E.** Expression of lipid catabolic genes *acs-2* and *cpt-3* is necessary for lifespan extension in *npp-16OE* animals. **F-G.** Fluorescence imaging reveals that *nhr-49* and *daf-16* activity are required for *acs-2p::GFP* induction in *npp-16OE* worms. Scale bar: 1 mm. n=4 independent experiments. **H-I.** *nhr-49* activity is required for the longevity phenotype of *npp-16OE* worms, whereas *daf-16* activity is not. **J.** RNAi knockdown of *nhr-49* blunts the mRNA induction of lipid catabolic genes in *npp-16OE* worms, whereas *hlh-30* RNAi partially rescues the induction of *lipl-2* and *lipl-3* in *npp-16OE* worms by qRT-PCR. n=3 independent experiments. **K.** Knockdown of *hlh-30* by RNAi inhibits lifespan extension in *npp-16OE* worms. Bars represent mean ± SD. Statistical significance was determined by two-way ANOVA (A, B, G, and J). Empty vector (EV) serves as the negative control for RNAi experiments. Relative mRNA levels in worm and HeLa cells were normalized to *act-1* (A and J) and Actin (B) respectively. ns: non-significant, \**p*<0.05, \*\**p*<0.01, \*\*\**p*<0.001, \*\*\*\**p*<0.0001. See also Table S1 for independent biological replicates and summary lifespan statistics.

Collectively, these data indicate that *npp-16OE* phenocopies the pro-longevity effects of hormetic starvation by mimicking the ability of energetic and nutrient stress to activate lipid catabolism. Since NPP-16/NUP50 is a nuclear basket protein without a known direct role in transcription, we next aimed to determine whether NPP-16/NUP50 recruits specific transcription factors to promote transcription of genes encoding lipid catabolic machinery. Several transcription factors have been reported to control *acs-2p::GFP* including potential activators NHR-49, DAF-16, HLH-30, ATFS-1, and PHA-4, and potential suppressor HLH-11^22,25,59,60^. An RNAi screen of these transcription factors revealed that only *nhr-49*/HNF4 and *daf-16/FOXO* RNAi are required for increased *acs-2p::GFP* reporter activity in *npp-16OE* worms (Figures 4F and 4G). Meanwhile, RNAi against *hlh-11* and *atfs-1* increases *acs-2p::GFP* reporter expression further in *npp-16OE* worms (Figures 4F and 4G), suggesting that regulation of *acs-2* by HLH-11 and ATFS-1 occurs in parallel to NPP-16. Consistent with a central role of *nhr-49/HNF4* in transcriptional regulation downstream of *npp-16,* RNAi knockdown of *nhr-49* also blunts extended lifespan and the increased mRNA level of lipid catabolic genes in *npp-16OE* worms (Figures 4H, 4J, and S4G).

Alternatively, *daf-16/FOXO* RNAi does not impact *npp-16OE* longevity (Figure 4I), suggesting that DAF-16/FOXO is not a dominant effector downstream of NPP-16. Although *hlh-30* knockdown by RNAi has no effect on the *acs-2p::GFP* reporter in *npp-16OE* worms (Figures 4F and 4G), *hlh-30* knockdown inhibits increases in lysosomal lipase gene expression and the corresponding lifespan extension of *npp-16OE* worms (Figures 4J and 4K). This is consistent with a known role for HLH-30/TFEB in governing transcriptional activation of lysosomal lipases upon starvation^57^. In aggregate, these results indicate that NHR-49 and HLH-30 are dominant transcription factors downstream of NPP-16/NUP50 that prompt activation of lipid catabolism and subsequent pro-longevity outcomes.

### Other nucleoporins and nuclear transport are dispensable for the metabolic and pro-longevity effects of NPP-16/NUP50

Given that NPP-16 has only minor effects on NPC permeability and assembly in worms^28,29^, we were curious about whether NPP-16 functions in concert with the NPC to modulate metabolism and aging. The nuclear basket of the NPC consists of NPP-21/TPR, NPP-16/NUP50, and NPP-7/NUP153, the latter serving as the anchor of NUP50/NPP-16 to the NPC^27^. By labeling the nuclear envelope with CRISPR-knock in of a mKate2 fluorescent tag to endogenous *npp-11* (NPP-11::mKate2), a nucleoporin localized in the central channel of the NPC, we found that post-developmental RNAi of *npp-7* expectedly prompted translocation of NPP-16 from the NE into the nucleoplasm (Figure 5A). Interestingly, neither *npp-7* nor *npp-21* RNAi have any effect on *acs-2p::GFP* reporter induction in *npp-16OE* worms, whereas *acs-2p::GFP* induction is mitigated by *npp-16* RNAi (Figure 5B). Consistent with these findings, the longevity phenotype of *npp-16OE* worms is only inhibited by post-developmental RNAi knockdown of *npp-16*, whereas it is unchanged by *npp-7* RNAi and *npp-21* RNAi (Figures 5C-5E). These data suggest that the interaction of NPP-16 with NPC is not essential for its metabolic and pro-longevity actions.

**Figure 5.**
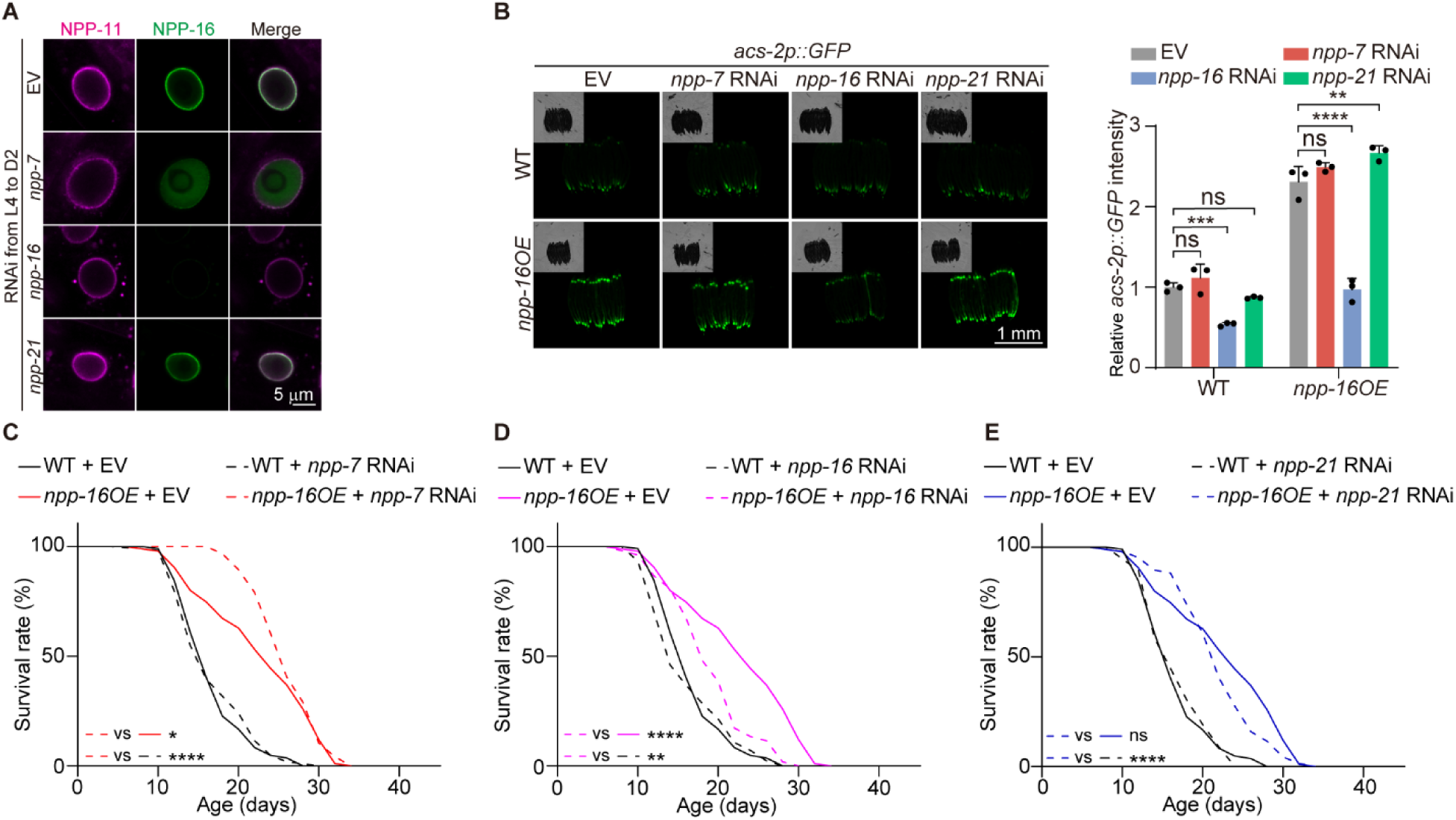
Other nucleoporins are not necessary for the pro-longevity effect of *npp-16OE*. **A.** Representative fluorescence images of endogenous *GFP::3xFLAG::AID::npp-16* and *npp-11::mKate2::3xFLAG::AID* abundance in the anterior intestinal cells of worms treated with the indicated RNAi from L4 to day 2 adulthood (D2). Scale bar: 5 μm. **B.** Knockdown of *npp-7* and *npp-21* by RNAi to D2 has no effect on the induction of *acs-2p::GFP* in *npp-16OE* worms. Knockdown was initiated at the late L4 stage of development to avoid developmental pleiotropy. Scale bar: 1 mm. n=3 independent experiments. **C-E.** Knockdown of *npp-7* and *npp-21* by RNAi has no effect on the lifespan extension seen in *npp-16OE* worms, which is expectedly suppressed by *npp-16* RNAi. RNAi was initiated at the L4 stage of development by a switch from empty vector (EV) to indicated RNAi. Bars represent mean ± SD. Statistical significance was determined by two-way ANOVA (B). Empty vector (EV) serves as the negative control for RNAi experiments. ns: non-significant, \**p*<0.05, \*\**p*<0.01, \*\*\**p*<0.001, \*\*\*\**p*<0.0001. See also Table S1 for independent biological replicates and summary lifespan statistics.

Although NPP-16/NUP50 is responsible for releasing cargo from importin-α during nuclear active transport^61^ and for NPC assembly during mitosis^62^ in mammalian cells, less is known about the roles of NPP-16/NUP50 in nuclear transport in post-mitotic cells, leading us to test whether NPP-16 promotes longevity by having an impact on nuclear transport. Speaking against this possibility, we find that *npp-16* knockdown does not impact nuclear localization of SV40 nuclear localization signal (NLS) tagged GFP, a canonical active nuclear transport cargo, whereas the knockdown of *ima-3* (orthologue of importin-α in the somatic tissues of *C. elegans*^63^) decreases the nuclear GFP signal significantly (Figure S5A). Consistent with the role of NPP-16 in longevity distinct from active nuclear transport, *npp-16OE* still extends lifespan significantly when *ima-3* is knocked down by RNAi (Figure S5B). In addition, knockdown of *ran-1*/RAN, which is the small GTPase essential for both active nuclear import and export^64,65^, is not epistatic to the longevity effects of *npp-16OE* (Figure S5C), indicating that *npp-16* extends lifespan in an active nuclear transport-independent manner. As the NPC barrier for passive diffusion is required for the pro-longevity and anti-cancer effects of metformin^20^, we also examined the effect of NPP-16 in maintenance of the passive transport barrier of the NPC. By quantifying the signal upon fluorescence recovery after photobleaching (FRAP) of a constitutively expressed intestinal GFP transgene, we find that *npp-21* knockdown by RNAi renders the nucleus more permeable to passive transport as reported^20^, whereas the recovery rate is unchanged by *npp-16* knockdown compared with empty vector control (Figure S5D).

Collectively, these results indicate that 1) NPP-16 is not required for active nuclear transport and NPC passive barrier maintenance in post-mitotic cells and 2) active nuclear transport and the passive NPC barrier are not required for lifespan extension in *npp-16* overexpression. Thus, we suggest a non-canonical role for NPP-16 in transcriptional modulation of metabolism and aging.

### The intrinsically disordered region of NPP-16/NUP50 is required for remodeling of the lipid catabolic transcriptome

As our evidence suggests that NPP-16/NUP50 regulates metabolism and longevity in a non-canonical manner independent of nuclear transport, we then hypothesized that NPP-16/NUP50 controls transcription directly via nucleoplasmic interactions. A significant feature of scaffold nucleoporins governing NPC permeability is the presence of phenylalanine-glycine (FG) repeat-containing, intrinsically disordered regions (IDRs), which mediate the interaction between protein NLSs and active nuclear transport machinery, and may also contribute to the passive diffusive barrier of NPC^66,67^. As a peripheral nucleoporin, NPP-16/NUP50 contains eight FG-repeats, six of which are enriched in the middle of the protein (Figure 6B), with no clearly established function. By bioinformatic prediction^68^, we identified a long IDR with a high propensity overlapping the central FG-repeat domain in NPP-16 (Figures 6A and 6B). Given that IDRs are also capable of mediating protein-protein and protein-DNA interactions and guiding transcription^69–72^, we hypothesized that the IDR in NPP-16 may be indispensable for its ability to modulate metabolism at a transcriptional level. To test this possibility, we constructed an integrated transgenic *C. elegans* strain overexpressing an NPP-16 variant without the central FG-repeats and IDR (lacking amino acids 244-354) driven by its native promoter (*npp-16*^Δ*IDR*^*OE*, Figure 6B). Importantly, this transgenic strain shares a similar expression level to our wild type *npp-16OE* (Figure 6C). Similar to NUP50 in mammalian cells^38^, NPP-16 colocalized with euchromatin and was excluded from heterochromatin marked by 4’,6-diamidino-2-phenylindole (DAPI) staining in *C. elegans*, whereas NPP-16^ΔIDR^ did not (Figure 6D), indicating that the IDR in NPP-16 regulates its association with chromatin regions with more open conformation.

**Figure 6.**
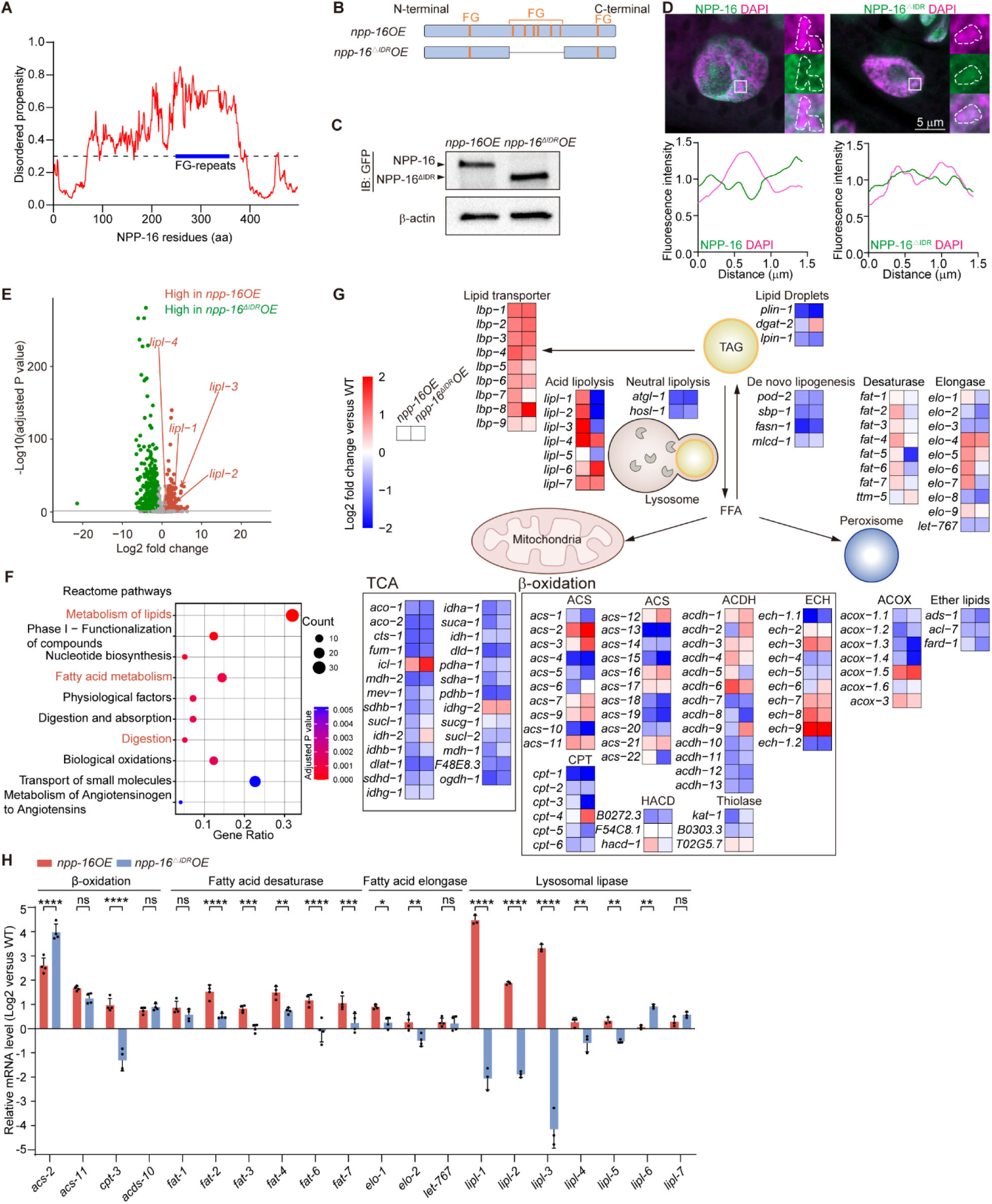
The intrinsically disordered region (IDR) of NPP-16/NUP50 is required for its ability to transcriptionally regulate lipid catabolism. **A.** NPP-16 harbors a predicted intrinsically disordered region (IDR) overlapping with its known FG-repeat sequences. The threshold predictive of an IDR of 0.3 is indicated by a black dashed line and the thick blue line indicates the FG-repeat domain. The IDR is predicted using flDPnn. **B.** Diagram displaying the structure of NPP-16 overexpression variants and the location of FG-repeats. **C.** Immunoblotting validates overexpression of full-length NPP-16 (*npp-16OE*) or a variant lacking the IDR (*npp-16*^Δ^*^IDR^OE*). β-actin serves as a loading control. **D.** Full-length GFP::NPP-16 does not overlap with condensed heterochromatin marked by DAPI staining, whereas GFP::NPP-16^ΔIDR^ does. The white dashed areas indicate the condensed chromatin and are 3.3x enlarged in the side panels. Scale bar: 5 μm. The relative fluorescence intensity was normalized to the mean absolute intensity of each region for quantification. **E.** Volcano plot highlighting differentially expressed genes (DEGs) from RNA-seq analyses of *npp-16OE* versus *npp-16*^Δ^*^IDR^OE* day 1 adult (D1) worms. Red and green dots represent the significantly up-regulated DEGs in *npp-16OE* and *npp-16*^Δ^*^IDR^OE* worms respectively, while gray dots are non-significantly altered genes. DEGs were determined using an adjusted *P* value < 0.05 and an absolute log_2_ fold change of 1. Significantly altered lipases are colored and marked with arrows as indicated. See also Table S3 for detailed statistics. **F.** Reactome pathway overrepresentation analyses of the up-regulated DEGs in *npp-16OE* worms versus *npp-16*^Δ^*^IDR^OE* D1 worms. Pathways related to lipid metabolism or lipid lipase activity are highlighted in red. See also Table S3 for detailed statistics. **G.** An integrated expression level and pathway diagram of lipid metabolic genes in *npp-16OE* and *npp-16*^Δ^*^IDR^OE* D1 worms versus WT in RNA-seq data. Values represent the log_2_ fold change in gene expression of npp*-16OE* (left column) or *npp-16*^Δ^*^IDR^OE* (right column) animals relative to WT. See also Table S3 for detailed statistics. TCA: tricarboxylic acid cycle. FFA: free fatty acid. **H.** qRT-PCR validation of lipid catabolic gene expression in *npp-16OE* and *npp-16*^Δ^*^IDR^OE* D1 worms. n≥3 independent experiments. Bars represent mean ± SD. Statistical significance was determined by two-way ANOVA (H). Relative mRNA levels were normalized to *act-1* (H). ns: non-significant, \**p*<0.05, \*\**p*<0.01, \*\*\**p*<0.001, \*\*\*\**p*<0.0001.

To define how the NPP-16/NUP50 IDR may regulate the lipid catabolic and metabolic adaptation transcriptome, we then performed bulk RNA sequencing (RNA-seq) in WT, *npp-16OE,* and *npp-16*^Δ*IDR*^*OE* worms at day 1 of adulthood. Using an adjusted *P* value threshold of 0.05 and absolute log_2_ fold-change of 1, we found that *npp-16OE* and *npp-16*^Δ*IDR*^*OE* shared 1069 up-regulated and 2459 down-regulated differentially expressed genes (DEGs) compared with WT, whereas *npp-16OE* worms have 947 up-regulated and 438 down-regulated DEGs that are IDR-dependent (Figure S6A). By GO-term analysis of the 947 IDR-dependent up-regulated DEGs^73,74^, we found enrichment for genes involved in lipid metabolism (Figure S6B and Table S3), consistent with our previous findings (Figures 1 and 4). To illustrate IDR-dependent transcriptional regulation, we also directly compared the transcriptomes of *npp-16OE* and *npp-16*^Δ*IDR*^*OE* animals (Figure 6E and Table S3). By Reactome pathway overrepresentation analysis and annotation^75,76^, we found that the pathways related to the metabolism of lipids, fatty acid metabolism, and lipid digestion are among the top 10 pathways enriched in the up-regulated DEGs of *npp-16OE* worms (Figure 6F and Table S3). By comparing the expression of metabolic genes relative to WT, we found that *npp-16OE* phenocopies the transcriptional alteration of starvation response^22,33,77^, in which the genes of tricarboxylic acid (TCA) cycle, lipid droplet biogenesis, and *de novo* lipogenesis are mostly inhibited. In contrast, lipid transporters are generally increased in *npp-16OE* worms mimicking the organismal starvation response^22^ (Figures 6G and 6H). While the genes involved in lipid β-oxidation are not globally increased or decreased, starvation-specific altered genes are regulated in a pattern similar to *npp-16OE*^22^, including *acs-2*, *acs-11*, and *hacd-1*, which are up-regulated; and *acdh-2*, *ech-1.1*, *ech-6*, *kat-1*, etc. which are down-regulated (Figure 6G). Strikingly, while *npp-16*^Δ*IDR*^*OE* regulates some metabolic genes similarly to *npp-16O*E, *lipl-1*, *lipl-2*, and *lipl-3* induction by *npp-16O*E is mitigated by loss of the IDR in *npp-16*^Δ*IDR*^*OE* worms (Figures 6G and 6H). In addition, most fatty acid desaturases that convert fatty acid into monounsaturated fatty acids (MUFA) and polyunsaturated fatty acids (PUFA) are up-regulated by *npp-16*, similar to the response to starvation^22^, but they are unchanged or decreased in *npp-16*^Δ*IDR*^*OE* worms (Figures 6G and 6H), suggesting that fatty acid composition is also altered in an IDR-dependent manner. Overall, these data suggest that lipid catabolic gene expression and emphatically the lysosomal acid lipases, are regulated by NPP-16/NUP50 in an IDR-dependent manner, raising the possibility that the IDR is also crucial for the pro-longevity outcomes associated with NPP-16/NUP50 overexpression.

### The IDR of NPP-16/NUP50 coordinates a longevity-promoting transcriptional response

We next explored the impact of the NPP-16/NUP50 IDR on the nucleoporin’s metabolic and pro-longevity effects of NPP-16/NUP50. As hypothesized, we found the lifespan of *npp-16*^Δ*IDR*^*OE* worms is similar to WT worms whereas the overexpression of full-length NPP-16 promotes longevity (Figures 4C and 7A). Neutral lipid fat mass is also unchanged in *npp-16*^Δ*IDR*^*OE* worms compared with WT worms (Figure 7B), showing that the IDR of NPP-16/NUP50 is necessary for the modulation of longevity and metabolism. As IDRs mediate protein-protein and DNA-protein interactions important for transcriptional regulation^69–72^, we next asked whether NPP-16/NUP50 directly associates with chromatin and the transcriptional machinery in an IDR-dependent manner. Immunoprecipitation of endogenous NPP-16 indicated direct binding to the activated transcriptional machinery *in vivo*, evidenced by interaction with phosphorylated RNA polymerase II^78^ (p-Pol II^Ser5^, Figure 7C). Endogenous NPP-16 displays enhanced recruitment onto the promoter of *acs-2* and *lipl-3* upon starvation by chromatin immunoprecipitation (ChIP, Figures 7D and S7A). Interestingly, ChIP qPCR in *npp-16OE* and *npp-16*^Δ*IDR*^*OE* worms reveals that the NPP-16/NUP50 variant lacking the IDR has no demonstrable interaction with the promoter of *acs-2* and *lipl-3* (Figures 7E and S7B), indicating that the IDR is required for the chromatin binding specificity of NPP-16/NUP50. Overexpression of *npp-16* is also sufficient to enhance the association of the transcriptional machinery (p-Pol II^Ser5^) with the promoter of *lipl-3* in an IDR-dependent manner (Figure 7F). Unexpectedly, we found that the *npp-16*^Δ*IDR*^*OE* promotes the engagement of active transcriptional machinery (through p-Pol II^Ser5^) on the promoter of *acs-2* (Figure S7C), which is consistent with its increased mRNA expression in *npp-16*^Δ*IDR*^*OE* worms (Figures 6G and 6H), suggesting that the transcriptional regulation of *acs-2* might be either an indirect effect from NPP-16, or a feedback output from elevated *lbp-8* expression, known to sufficiently promote *acs-2* expression and longevity^79^. Overall, these data demonstrate that NPP-16/NUP50 is necessary and sufficient for the recruitment of transcriptional machinery onto the *lipl-3* promoter in a manner dependent on the presence of the IDR.

**Figure 7.**
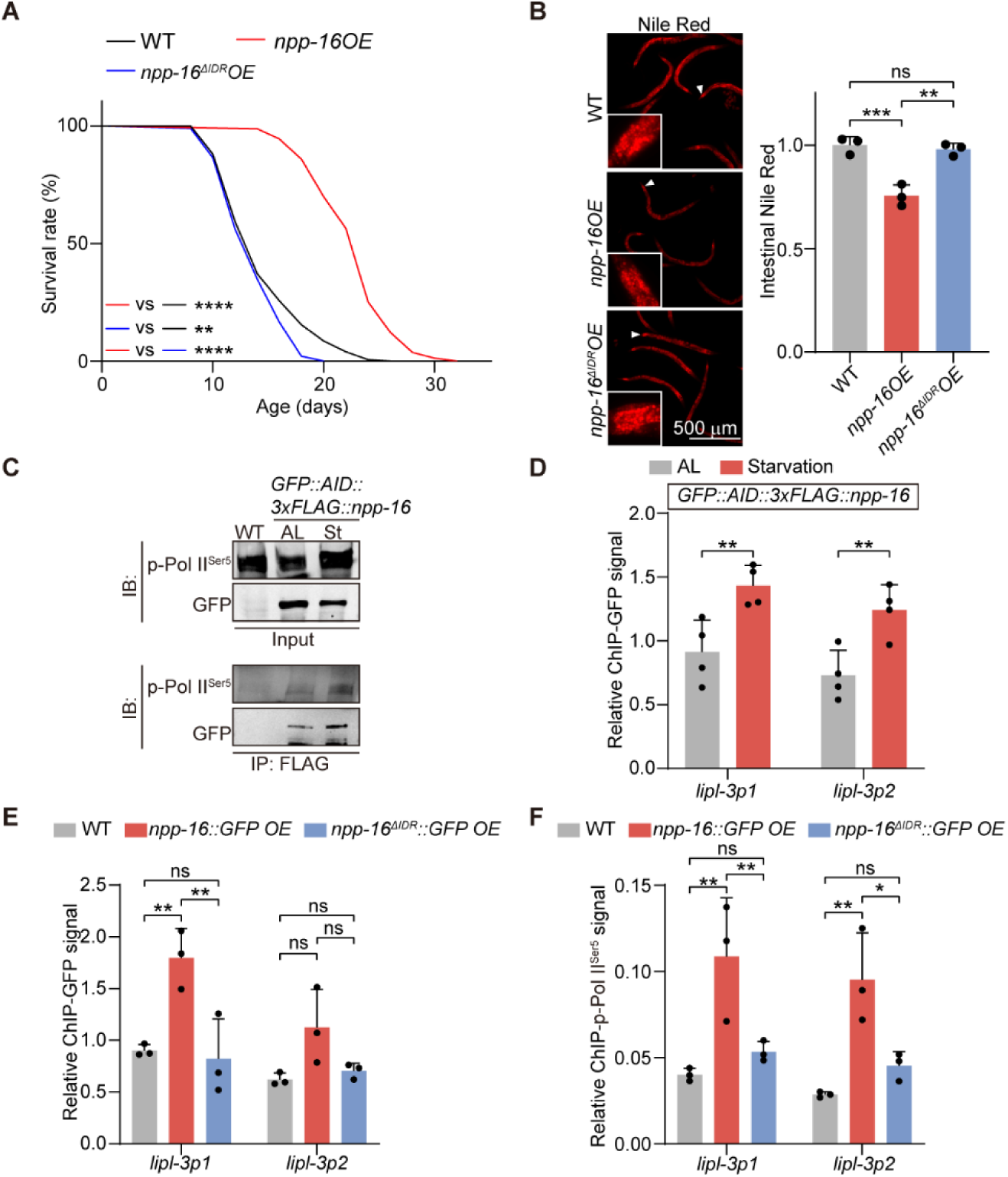
The IDR is required for the metabolic and pro-longevity actions of NPP-16/NUP50. **A.** Lifespan analyses reveal that transgenic *C. elegans* overexpressing a NPP-16 variant lacking the intrinsically disordered region (*npp-16*^Δ^*^IDR^OE*) no longer exhibit lifespan extension evident in *npp-16OE*. **B.** The IDR is required for *npp-16OE*-mediated fat mass reduction in the posterior intestinal cells, which are indicated by white arrowheads and enlarged in white boxes. Scale bar: 500 μm. n=3 independent experiments containing at least 60 worms per group. **C.** Endogenous GFP::3xFLAG::AID::NPP-16 binds to phosphorylated RNA polymerase II (p-Pol II^Ser5^) by co-immunoprecipitation. **D.** Starvation for 4 hours enhances the association of endogenous GFP::AID::3xFLAG::NPP-16 with the *lipl-3* promoter. n=4 independent experiments. *lipl-3p1* and *lipl-3p2* are two distinct regions within the *lipl-3* promoter. **E.** ChIP-qPCR reveals that full-length NPP-16 in *npp-16OE* worms binds to the *lipl-3* promoter *in vivo*, whereas the NPP-16^ΔIDR^ in *npp-16*^Δ^*^IDR^OE* worms does not. n=3 independent experiments. **F.** ChIP-qPCR reveals that *npp-16OE* but not npp*-16*^Δ^*^IDR^OE* promotes the binding of p-Pol II^Ser5^ to the *lipl-3* promoter. n=3 independent experiments. Bars represent mean ± SD. Statistical significance was determined by one-way ANOVA (B) and two-way ANOVA (D-F). Relative ChIP signals were normalized to *act-1* promoter (D-F). See also Table S1 for independent biological replicates and summary lifespan statistics. ns: non-significant, \**p*<0.05, \*\**p*<0.01, \*\*\**p*<0.001, \*\*\**p*<0.0001.

## Discussion

Nucleoporins are well-known for mediating nucleocytoplasmic transport, a process crucial for transcriptional regulation and genome integrity^11^. The barrier functions of the NPC are disrupted with age and in certain neurodegenerative diseases, at least in part through deterioration of scaffold nucleoporins^14–16^. In spite of these known roles of the NPC, in this manuscript, we report a heretofore unappreciated mechanism by which NPC impacts metabolism and lifespan beyond nuclear transport. Specifically, we identify a peripheral nucleoporin in the nuclear basket which relays information from the energy sensor AMPK to the transcriptional machinery. Direct phosphorylation of NPP-16/NUP50 by AMPK is both necessary and sufficient to promote transcriptional adaption and the deployment of metabolic stress defenses in aging. Multiple, energetic longevity paradigms leverage this mechanism, which illuminates a new dimension of nucleoporin function in aging beyond active and passive nuclear transport (Figure S7D).

### A noncanonical role of the nuclear pore in metabolism and modulation of aging

Although the NPC can mediate transcription *in situ*^80,81^, the physiological functions and precise molecular mechanisms of this non-transport-linked action, and their potential relevance to aging are incompletely understood. As a dynamic nucleoporin, NUP50 was previously reported to localize into euchromatin upon heat stress in *Drosophila*^81^, implying a direct role of NUP50 in transcriptional regulation. In this study, we uncovered a new role for NPP-16/NUP50 in the positive transactivation of lipid catabolic gene transcription upon nutrient stress. This action is independent of the localization of NUP50 within the NPC and NUP50-binding nucleoporins, indicating a heretofore unappreciated role of NPP-16/NUP50 on chromatin accessibility. This function is consistent with NUP50 having the highest mobility amongst all nucleoporins^82,83^, and that mobility depends upon normal transcription^38^. Most nucleoporins are stable with a long half-life associated with their scaffold role within the NPC, with any perturbations resulting in altered permeability of the nuclear envelope, compromising nucleocytoplasmic fidelity and organismal survival^29^. In HeLa cells, NUP50 regulates active nuclear transport by releasing transport cargoes from importin-α and mediates nuclear assembly during cell division^27,62^. Genomic deletion of NUP50 also causes late embryonic lethality in mice^84^, however, its disruption does not affect fibroblast or myoblast proliferation^38,84^, suggesting developmental timing- or cell-lineage-specific functions of NUP50 in nuclear pore transport and assembly. In worms, our data and the data of others indicate that NPP-16/NUP50 perturbation has no obvious impact on the NPC barrier, nuclear transport, larval development, or lifespan of wild-type *C. elegans*^28,29^, indicating that the function of NPP-16 in canonical NPC assembly and nucleocytoplasmic barrier integrity might be less important in nematodes.

Our data point very specifically to the modulation of NUP50. In this study, we find that nucleoporin protein induction responding to nutrient and energetic stress is specific to NUP50, and that other scaffold nucleoporins are unchanged. This unique induction of NUP50 at both the NE and nucleoplasm implies the binding ratio of NUP153 and NUP50 at NPC is flexible and affected by energetic status. Since NUP153 has two distinct interacting sites with NUP50 and its protein level is unchanged by nutrient stress^27^, nutrient availability might determine the relative availability of NUP50 to participate in metabolic rewiring. The specific induction of NUP50 also suggests dynamic modulation of NPC composition and structure upon physiological stimuli *in vivo*. While technically challenging to assess, we suggest that the biophysical and binding properties of NUP50 at the NPC, its shuttling within the nucleoplasm, and the general stoichiometry of nucleoporins at the NPC under multiple stressors could be of great interest for further exploration.

### Connecting energy sensing to metabolic adaptation and longevity

Our findings are the first to implicate NUP50 as a link between energetic status, AMPK, and pathways that alter energy production. We suggest that NUP50 is a previously unappreciated cornerstone of metabolic adaptation in aging. AMPK is a well-established hub for energy sensing and metabolic adaptation upon nutrient and energetic stress, activated by increased LKB1-mediated phosphorylation in the face of a rising AMP/ATP ratio, for example with biguanide treatment^1,41^. AMPK has well-described responses to energetic stress, including direct phosphorylation and inhibition of acetyl-CoA carboxylase (ACC), which leads to reduced malonyl-CoA production, allosteric activation of carnitine palmitoyl transferase (CPT1), and ultimately enhanced lipid β-oxidation for energy production^85^. Previous studies have highlighted parallel transcriptional metabolic shifts caused by activated AMPK, which is controlled by the orchestration of master transcriptional regulators CRTC-1/CRTC and NHR-49/HNF4^86^. Here we identify NUP50/NPP-16 as a new metabolic switch required for the response to nutrient status and increased AMPK activity, modulating lipid catabolism and metabolic adaptation in a manner dependent on NHR-49/HNF4 and HLH-30/TFEB. These findings imply that this nuclear AMPK signaling cascade could be a vital part of the checks and balances that establish proper metabolic regulation and longevity in response to energetic status. Thus, we put forward the mitochondria to nucleus AMPK/NUP50 signaling axis as an essential bridge between energy sensing and metabolic adaptations. Additionally, since most identified phosphorylation targets of AMPK occur in proteins that localize to the cytosol, mitochondria, or on lysosomal membranes^1^, the data we present highlight the importance of further study of nuclear-localized substrates of AMPK. Determining organelle-specific substrates of this master energy sensor is critical to understand its physiological and pathological functions in aging.

Hormesis is a benefit provided by a mild environmental challenge followed by the activation of stress defenses and positive effects on healthy aging. Nutrient stresses such as dietary restriction, exercise, and mild oxidative stress promote hormesis across species^87,88^. Many different regimes of nutrient and caloric deprivation are sufficient to extend lifespan in multiple species, such as intermittent fasting, global dietary restriction, and selective nutrient deprivation, resulting in an organismal metabolic switch to an adaptive ‘low energy’ mode by activating lipid catabolism and autophagy^44–47^. Here, we affirm prior observations that short-term food deprivation in early adulthood is sufficient to extend the lifespan of *C. elegans*, in the process placing NPP-16/NUP50 on the map of critical effectors of metabolic hormesis in aging. The fact that NPP-16/NUP50 is required during the narrow window of hermetic starvation, but not thereafter, has several important implications. First, NPP-16/NUP50 is critical to generating a proper initial response to nutrient stress, and its presence during the starvation phase is required for long-term benefit. Second, and curiously, NPP-16/NUP50 is not part of the machinery required to maintain the beneficial effects of hormetic starvation, invoking a discreet set of genes for the latter effect. These data imply that other genetic regulators may be involved in enforcing the beneficial effects of hormetic starvation after the initial impact of NPP-16/NUP50, such as epigenetic rewriters for lateral effect maintenance. Finally, we do not know whether the dramatic lifespan extension seen with overexpression of NPP-16 leverages additional benefits beyond those programmed by hormetic starvation. We are conclusively able to say that NPP-16 promotes longevity in a manner dependent upon genes of lipid catabolism, irrespective of whether additional upstream mechanisms are invoked.

### Roles of intrinsically disordered regions in aging

Our data suggest that NUP50’s IDR is critical for extending survival in response to nutrient stress. Given the transcriptomic differences between *npp-16OE* and *npp-16*^Δ*IDR*^*OE* worms, we identified lysosomal lipases and fatty acid desaturases as the dominant, specific transcriptional effectors downstream of the IDR in NPP-16/NUP50, which are activated similarly upon starvation^22^. Our data identify the essential cadre of IDR-dependent genes that drive longevity in response to nutrient stress. Nonetheless, and perhaps more surprisingly, some genes are similarly regulated in the presence or absence of the NPP-16/NUP50 IDR. Explicitly, some lipid catabolic genes, such as *acs-2* and *lbp-8*, whose overexpression sufficiently drives longevity in worms^26,79^, are also increased in an IDR-independent manner; nonetheless, their induction in *npp-16*^Δ*IDR*^*OE* worms is not sufficient to prompt lifespan extension. We posit that this is the result of the reduction of other critical metabolic genes limiting the pro-longevity benefits of *acs-2* and *lbp-8* activation, such as *lipl-3* and *cpt-3*, controlling the initiation of lipolysis and serving as the rate-limiting enzymes of fatty acid oxidation (FAO), respectively ^58^. Moreover, we also identified a group of IDR-independent *npp-16*OE DEGs, which may function as a separate regulon from our study that buffers the positive effects of sustained *acs-2* and *lbp-8* expression.

Interactions between transcription factors, co-transcriptional machinery, promoter architecture, and other cis-regulatory elements determine proper transcriptional activity. IDRs are a group of proteins with uncertain structure, potentially mediating protein/protein and protein/DNA interactions, which are critical for target recognition in transcriptional factors^69–72^. Recently, FG-repeat enriched nucleoporins have been reported to regulate NPC permeability through their IDRs. The IDRs are also critical for the capability of phase transition in these nucleoporins for mediating nuclear transport and permeability^89,90^. However, our study reveals a new action of the IDR in a nuclear basket protein that serves directly as a transcriptional cofactor, coordinating active transcriptional machinery on metabolic genes upon nutrient and energetic stress. The observed collaboration between the NPC and the transcriptional machinery is an underappreciated dimension of the response to environmental challenges. We also found that the IDR determines NPP-16/NUP50 localization in the nucleoplasm and specifically its association with open versus closed chromatin. Since chromatin reorganization has been shown to be promoted by mitochondrial stress for pro-longevity outcomes^91^, we speculate that an additional role of the NUP50 IDR could be to regulate transcription through modulation of chromatin accessibility and related epigenetic modifications.

### Conserved action of NUP50 in metabolic adaptation in mammals

Compellingly, we find that critical structural and regulatory elements of NUP50 important to aging are conserved from nematode to human. First, NPP-16/NUP50 structure, including its IDR and FG-repeat rich domains are conserved across species. Second, several pieces of evidence suggest the potential roles of NUP50 in age-related phenotypes. NUP50 dynamics in the nucleoplasm have been shown to be associated with global transcriptional activity and myoblast differentiation in mammalian models^38^. A recent study identified NUP50 as a high-risk gene in amyotrophic lateral sclerosis (ALS) patients^92^. Our study finds that the AMPK consensus phosphorylation motif and IDR in NPP-16/NPP-50 is conserved between *C. elegans*, mouse, and human, suggesting that NPP-16/NUP50 activation upstream of lipid catabolism in a manner independent from NPC activity may be an ancient and conserved function of the nucleoporin.

Parallel increases in human NUP50 levels during nutrient stress and elevated lipid catabolic gene expression by NUP50 overexpression in HeLa cells prompt us to speculate that metabolic and pro-longevity actions of NUP50 could also be conserved in higher eukaryotes. Further work will be required to define the full spectrum of NUP50 roles in aging and metabolism in order to put forward new therapeutic strategies to reduce age-related morbidity in humans.

## STAR Methods

### Key resource table

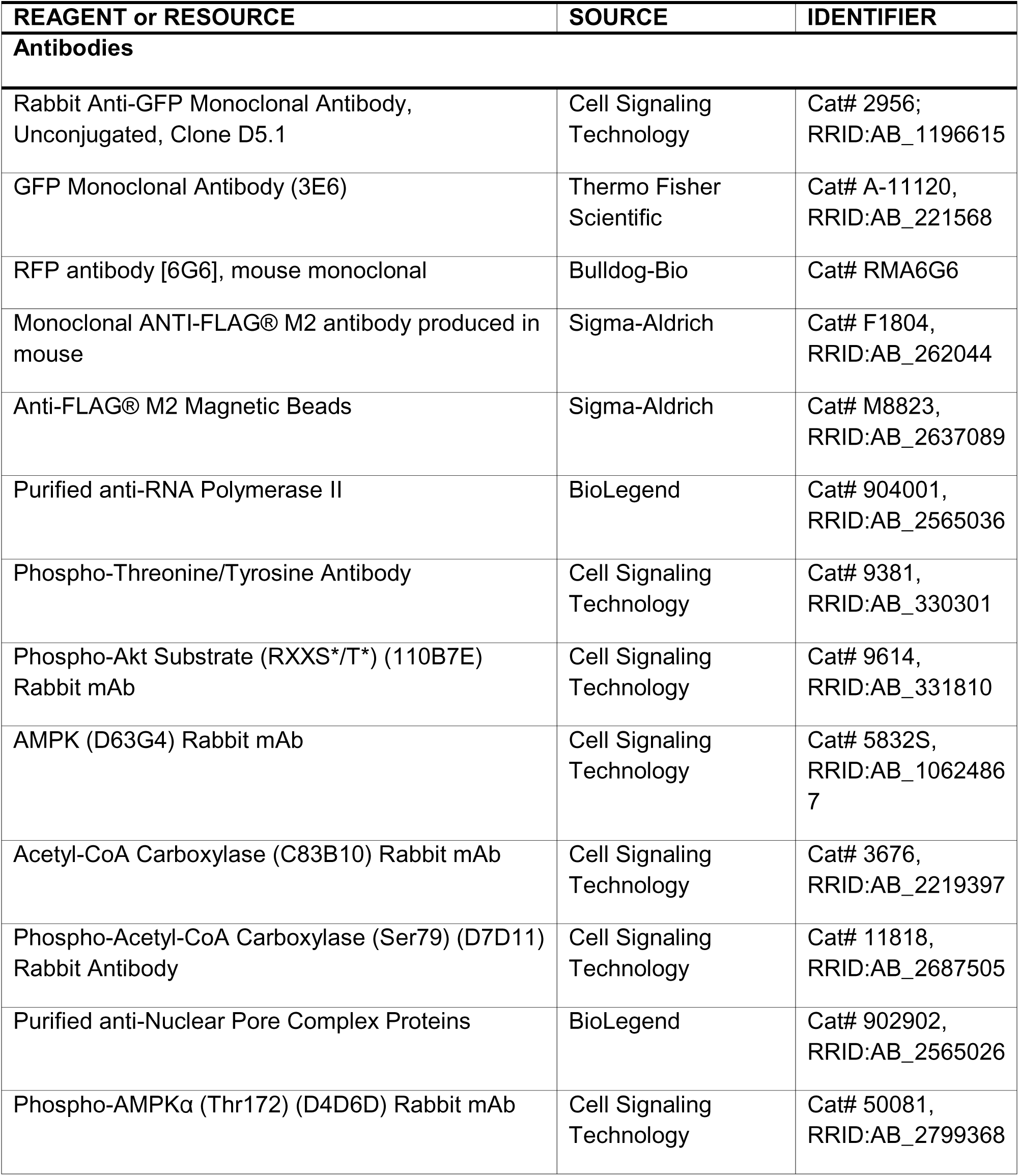
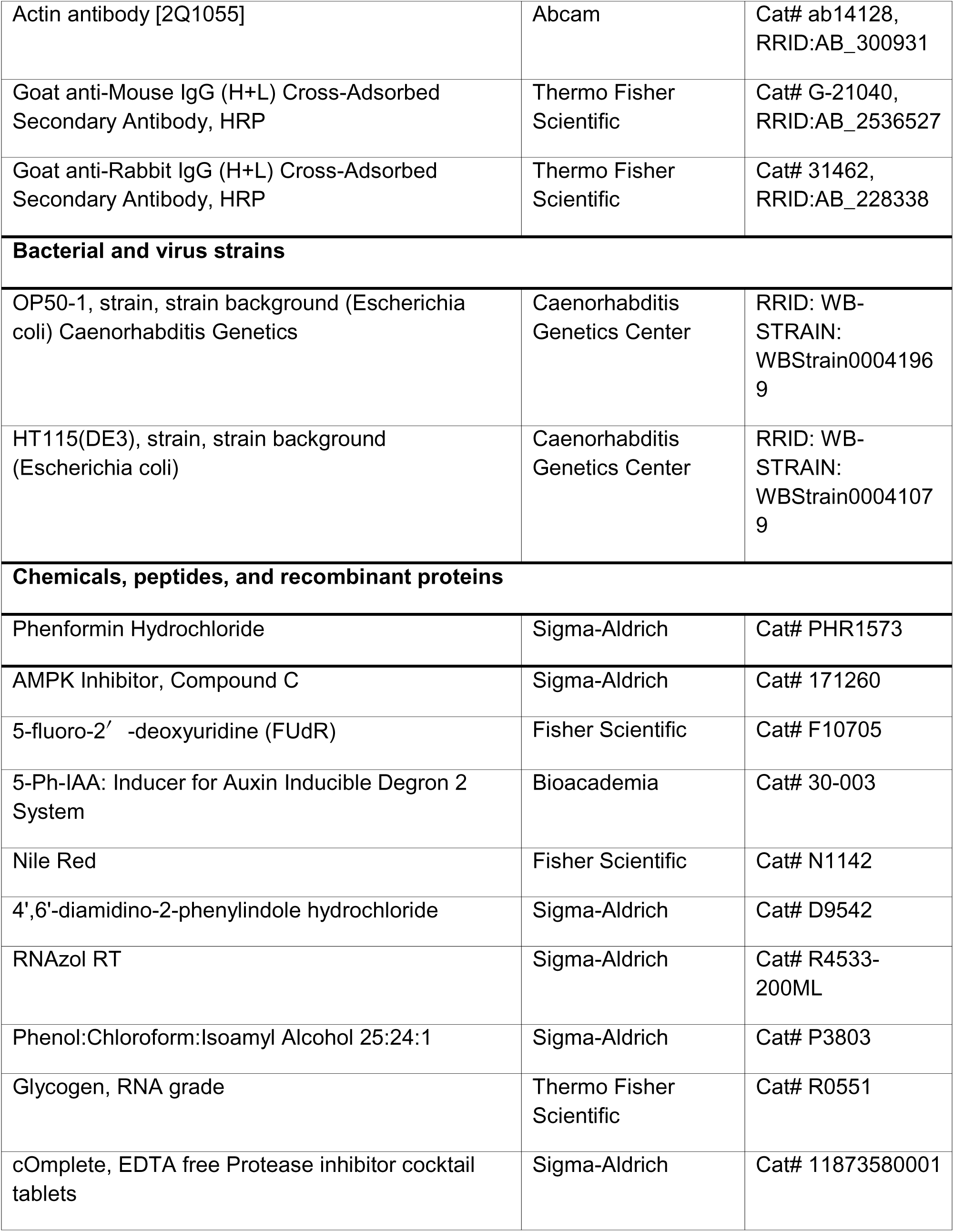
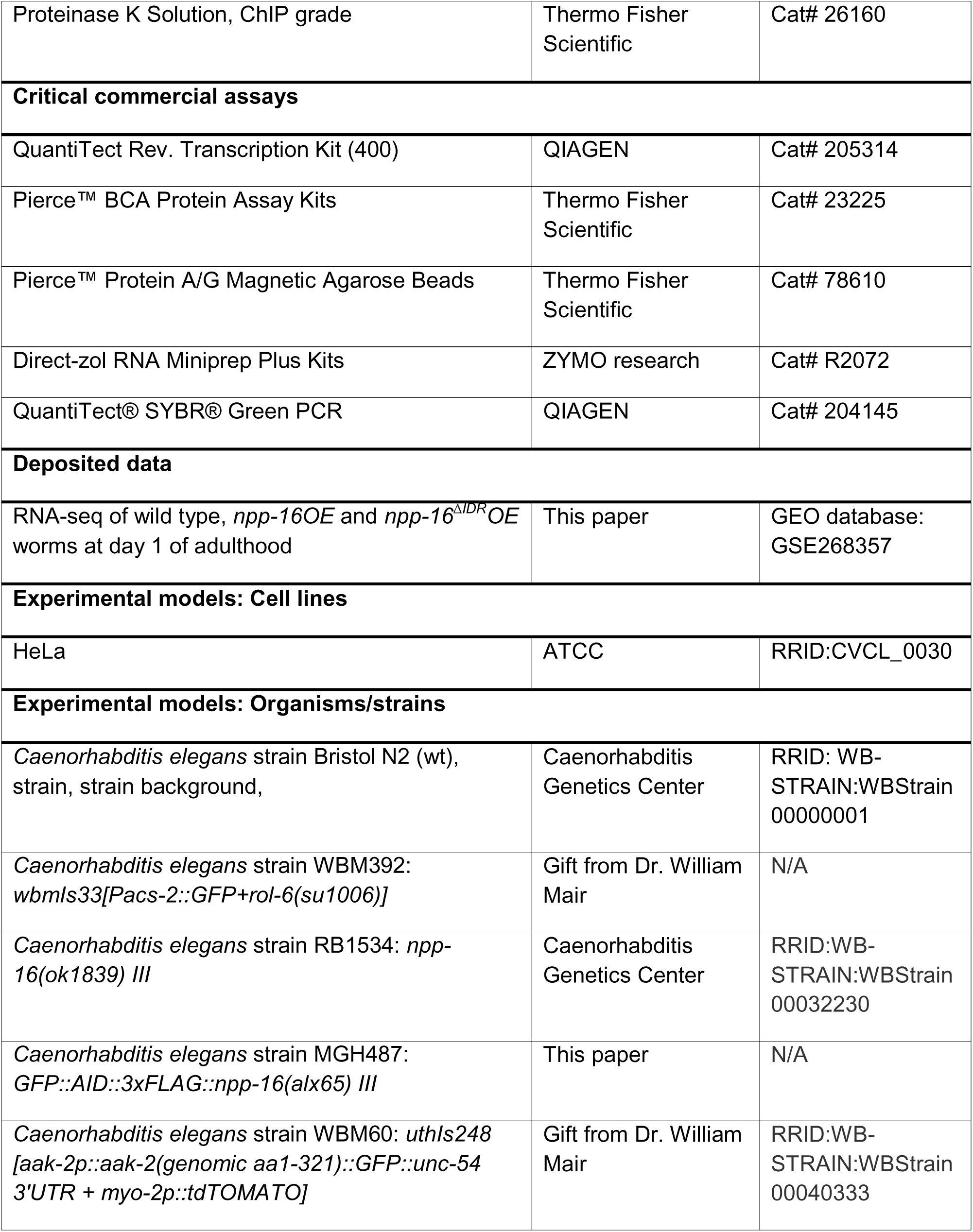
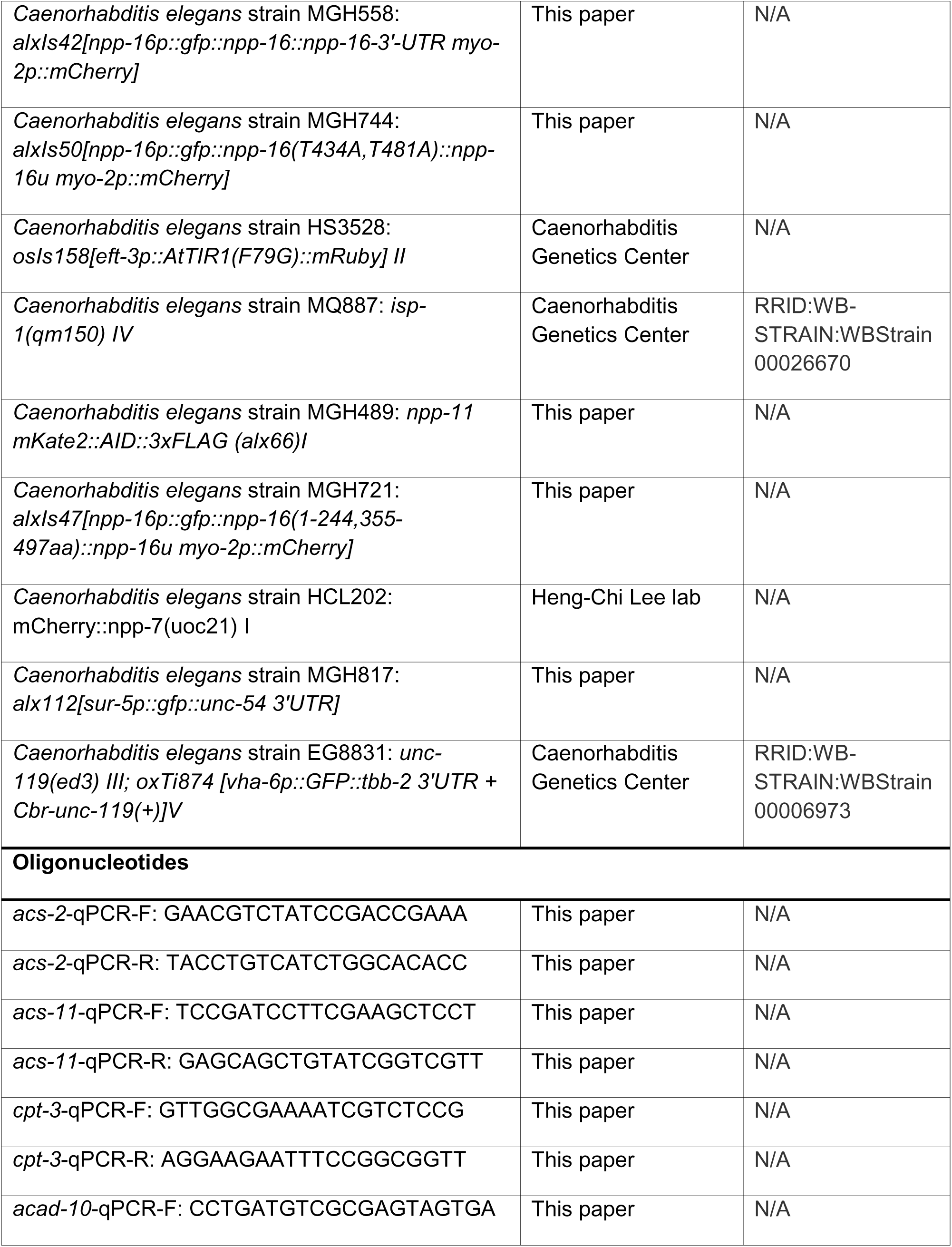
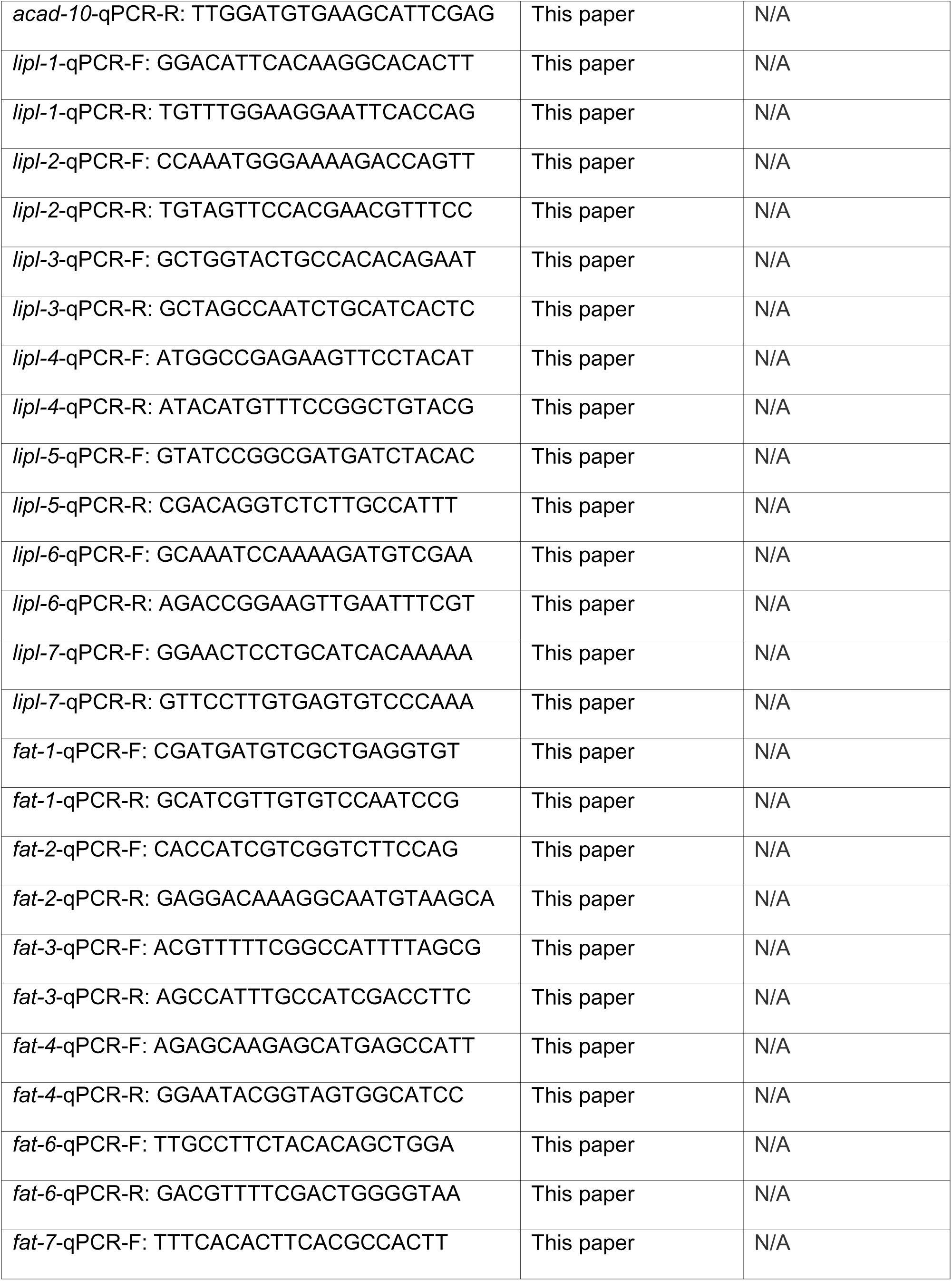
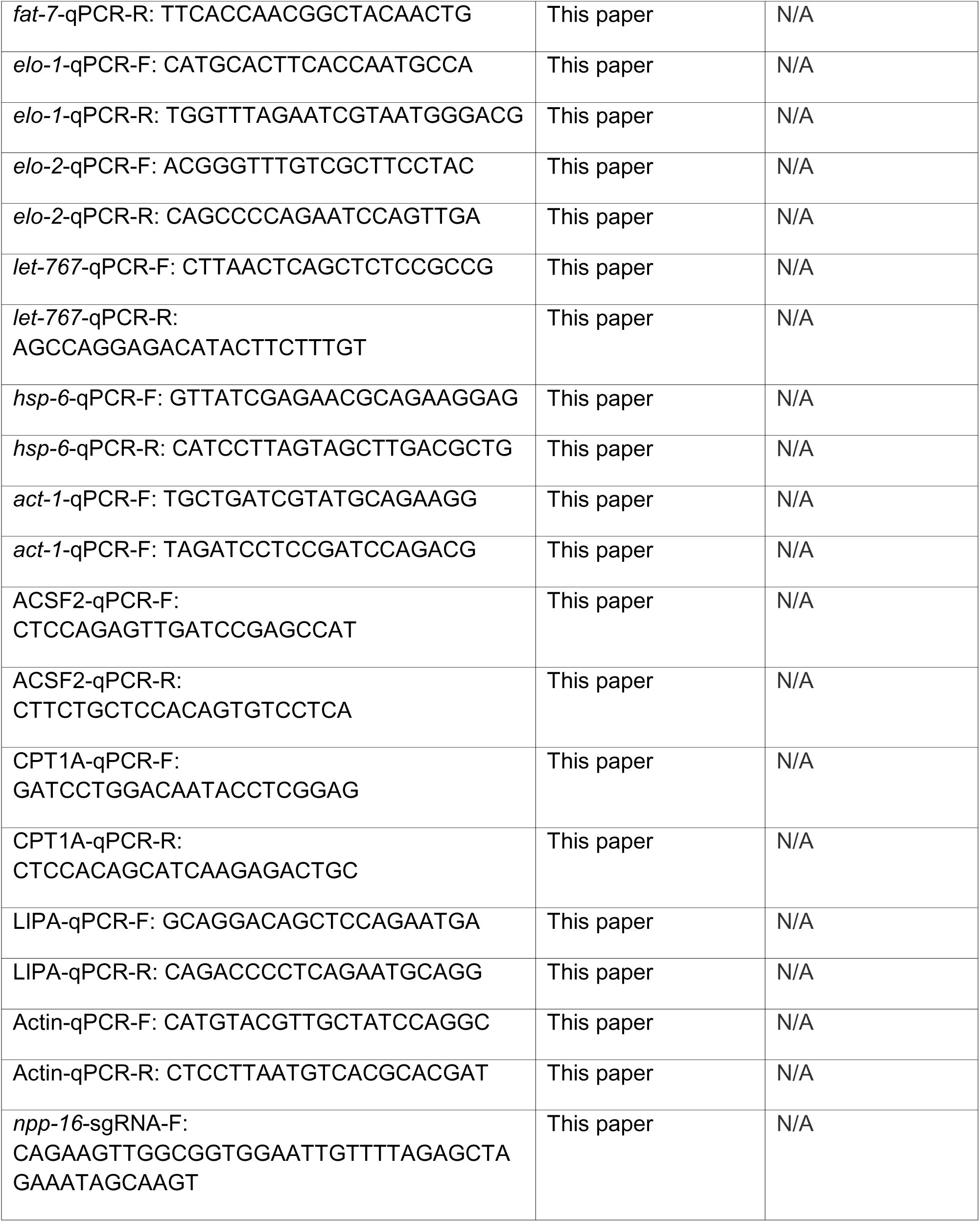
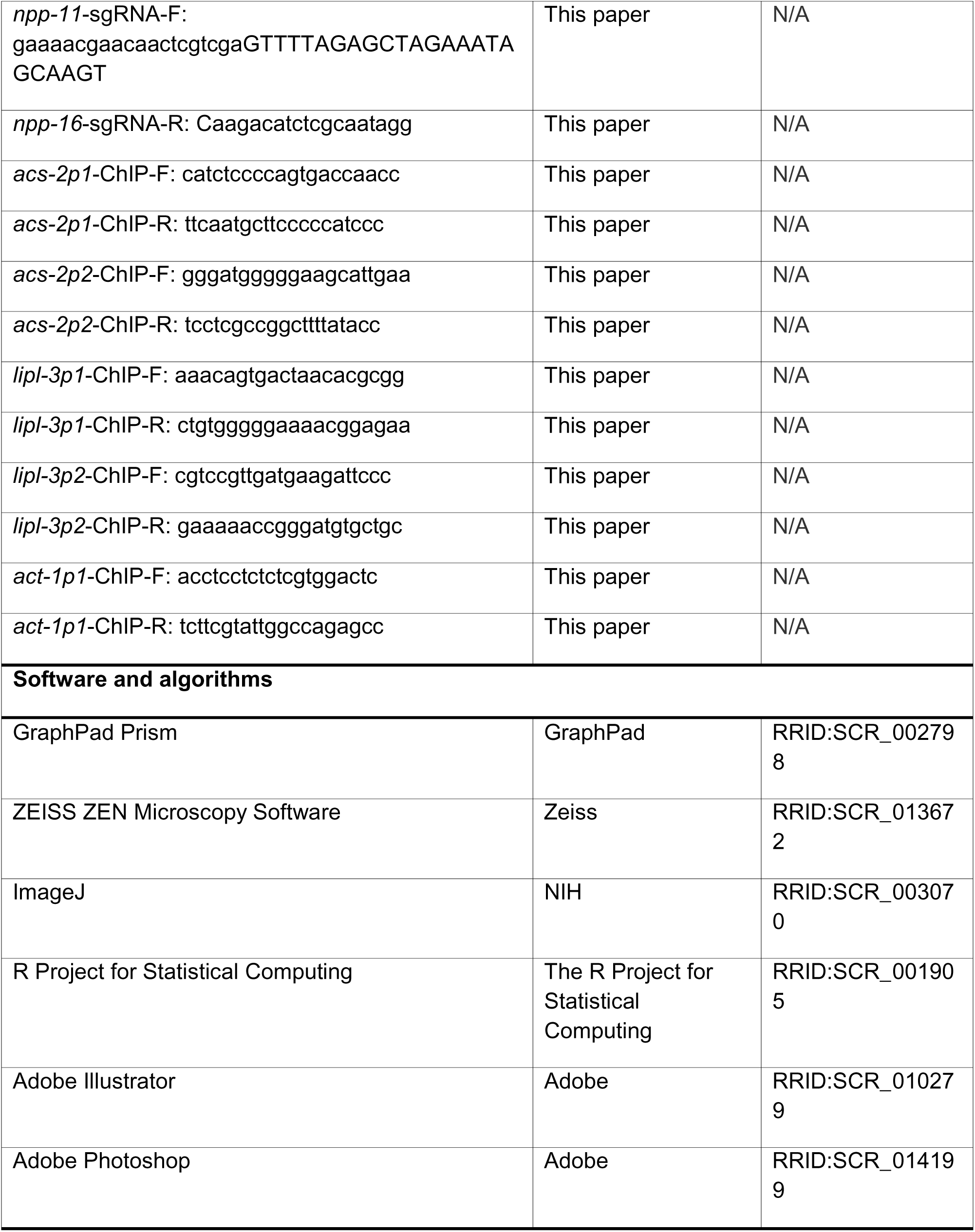

### Resource availability

#### Lead contact

Further information and requests for resources and reagents should be directed to and will be fulfilled by the lead contact, Alexander A Soukas.

#### Material availability

*C. elegans* strains and plasmids used in this study are available from the lead contact.

#### Data and code availability

This paper does not create any original code. The RNA-seq data have been deposited to the GEO database and assigned the accession number ‘GSE268357’.

### Experimental model and study participant details

#### C. elegans

*C. elegans* were grown on the normal growth medium (NGM) supplemented with 0.1875 mg/ml streptomycin and fed with the E.coli strain OP50-1 or HT115(DE3) with indicated RNAi under the standard condition at 20 °C unless otherwise noted^93^. All the strains used in this paper are listed in the key resource table; some of them were provided by the Caenorhabditis Genetics Center funded by the NIH Office of Research Infrastructure Programs (P40 OD010440), and some were kindly provided by the laboratory of William Mair at the Harvard T.H. Chan School of Public Health and the Heng-Chi Lee lab at University of Chicago.

For auxin treatment, the auxin analog 5-Ph-IAA (Bioacademia, Cat#30-003) was dissolved in DMSO at a stock concentration of 100 mM and diluted to 100 μM in water as a working solution immediately before use. Thereafter, 450 μl of the working solution was added onto the 9 mL of NGM in a 6 cm petri dish seeded with OP50 or HT115 bacteria, leading to a final concentration of 5 μM in agar. Auxin was added to plates one day before plate use, and plates were kept in the dark. The DMSO of the same dilution rate serves as the negative control.

For phenformin treatment, phenformin hydrochloride (Sigma-Aldrich, Cat# PHR1573) was dissolved in water at 0.1 M as a stock solution, and 425 μL of the stock solution was added onto 9 mL of NGM agar in a 6 cm petri dish seeded with OP50 or HT115 bacteria, leading to a final concentration of 4.5 mM. Phenformin was added one day before plate use.

#### Cell lines

HeLa cells were from ATCC and cultured in DMEM supplemented with 10% fetal bovine serum (Thermo Fisher Scientific, Cat# A3382001) at 37°C, 5% CO_2_. For serum starvation, the medium was replaced by DMEM without FBS washing cells gently several times with PBS, when the cell density was about 80%. The cells were collected for further experiments after being starved for the times indicated in figure legends. For AMPK inhibition, Compound C (Sigma-Aldrich, Cat# 171260) was dissolved in DMSO to a concentration of 100 mM as a stock solution and diluted to 20 μM in cell medium immediately prior to conducting experiments. For NUP50 overexpression, 2 μg of each plasmid was transfected into cells in each well of 6-well plates with Lipofectamine™ 3000 Reagent (Thermo Fisher Scientific, Cat# L30000015) following the manufacturer’s instructions.

## Method details

### Plasmid construction

For genome editing by CRISPR/Cas9, plasmids were constructed as reported^36,94,95^. To generate pJW1583-*GFP::AID::3xFLAG::npp-16* as the repair template for *npp-16* genomic editing, 519 bp of upstream and 525 bp of downstream sequence flanking the start codon of genomic *npp-16* was cloned into pJW1583 digested by SpeI and ClaI using Gibson assembly. To generate pJW1586-*npp-11::mKate2::AID::3xFLAG* as the repair template for *npp-11* genomic editing, 581 bp of upstream and 586 bp downstream sequence flanking the stop codon of genomic *npp-11* was cloned into the pJW1586 vector digested by SpeI and AvrII through Gibson assembly. The pJW1583 and pJW1586 vectors were provided by Addgene (Cat#121054 and Cat#121057).

To generate plasmids expressing guide RNAs for CRISPR editing, pDD162-*npp-16* and pDD162-*npp-11*, the sgRNA sequences were designed by CRISPOR (http://crispor.gi.ucsc.edu/crispor.py)^96^. The guide sequences (cagaagttggcggtggaatt and gaaaacgaacaactcgtcga) targeting the N-terminus of genomic *npp-16* and C-terminus of *npp-11* respectively were cloned into pDD162 by PCR and T4 DN ligase (New England Biolabs, Cat#M0202S). The pDD162 vector was provided by Addgene (Cat#47549).

To generate pDD95.79-*npp-16p::gfp::npp-16::npp-16u*, the genomic *npp-16* sequence containing a flexible linker (GGGGS, 1837 bp) and its 3’-UTR (117 bp) were cloned into the pDD95.79 vector linearized at the stop codon of GFP to generate pDD95.79*::gfp::npp-16::npp-16u*. The *npp-16* promoter (3428 bp) was then cloned into the subclone digested by XmaI through Gibson assembly.

To generate pDD95.79-*npp-16p::gfp::npp-16(T434/481A)::npp-16u*, site directed mutagenesis was performed on pDD95.79-*npp-16p::gfp::npp-16::npp-16u* by PCR with the primers carrying the T434A mutation using the QuikChange Lightning protocol. After validation with Sanger sequencing, T481A mutagenesis was performed with the same strategy on pDD95.79-*npp-16p::gfp::npp-16(T434A)::npp-16u*.

To generate pDD95.79-*npp-16p::gfp::npp-16*^Δ*IDR*^*::npp-16u*, a flexible linker (GGGGS), the split cDNA sequences of *npp-16* without the IDR (732 bp and 432 bp), and its 3’-UTR (117 bp) were cloned into the pDD95.79 vector linearized at the GFP stop codon by Gibson assembly. The *npp-16* promoter (3428 bp) was then cloned into the resulting subclone by Gibson assembly after XmaI digestion.

To generate pDNA3.1-FLAG-NUP50-GFP, the human NUP50 cDNA (1403 bp) with an N-terminal FLAG tag was amplified by PCR and cloned into the pcDNA3.1-EGFP vector digested by EcoRI and NotI through Gibson assembly.

### *C. elegans* strain generation

For the genome editing of *npp-16* by CRISPR/Cas9, plasmids of pDD162-*npp-16*(10 ng/μl) and pJW1583-*GFP::AID::3xFLAG::npp-16* (50 ng/μl) were co-injected into N2 worms with an injection marker of *myo-2::mCherry* (2.5 ng/μl). Similarly, for genome editing on *npp-11*, plasmids of pDD162-*npp-11* (10 ng/μl) and pJW1586-*npp-11::mKate2::AID::3xFLAG* (50 ng/μl) were co-injected into N2 worms with an injection marker of *myo-2::mCherry* (2.5 ng/μl). Salmon sperm DNA was added as a carrier to bring the injection mix final concentration to 100 ng/μl of DNA. The homozygotes with single copy knock-in were screened with hygromycin resistance assay as reported^36^.

To generate the integrated strain by UV radiation for overexpressing *npp-16* variants, the extrachromosomal arrays were generated first by injecting pDD95.79-*npp-16p::gfp::npp-16::npp-16u* (50 ng/μl), pDD95.79-*npp-16p::gfp::npp-16(T434/481A)::npp-16u* (50 ng/μl), and pDD95.79-*npp-16p::gfp::npp-16*^Δ*IDR*^*::npp-16u* (50 ng/μl) into N2 worms for *npp-16OE*, *npp-16(T434/481A)OE* and *npp-16*^Δ*IDR*^*OE* transgenic strains, respectively. *myo-2p::mCherry* (2.5 ng/μl) served as a co-injection marker and salmon sperm DNA was used as a carrier to bring the injection mix final concentration to 100 ng/μl of DNA. For transgene integration, the extrachromosomal arrays were radiated by exposure to 225 mJ and 250 mJ of UV irradiation at late L4, and the homozygotes with integrated transgenes were screened out by examining clonal populations of F2 or F3 progeny for homozygous red fluorescence in the pharynx.

### Lifespan assay

Worms were synchronized by overnight egg lay. After 60 hours, ∼30 late L4s were picked onto the NGM agar 6 cm plates seeded with OP50 or HT115 containing 50 mM 5-fluoro-2′-deoxyuridine (FUdR). Worms were counted every other day, and the ones not exhibiting spontaneous movement or subsequently not responding to mechanical prodding were scored as dead. Animals that exhibited bursting vulva or plates that became contaminated were censored. Statistical analysis was performed using the Mantel-Cox Log Rank in GraphPad Prism. See also Table S1 for independent biological replicates and summary lifespan statistics.

### RNA interference

RNAi clones were isolated and sequence validated from the Ahringer library^97^. The RNAi plates were made with the standard NGM recipe supplemented with 5 mM IPTG and 200 μg/ml carbenicillin. After overnight culture in LB containing 200 μg/ml carbenicillin at 37 °C, the bacteria were concentrated 5x by centrifuge. 200 μl concentrated bacteria were seeded onto the RNAi plates for corresponding experiments. All RNAi experiments were conducted from hatching unless otherwise noted.

For the RNAi screen on the *acs-2p::GFP* reporter with phenformin treatment, WBM392 worms were synchronized by egg-laying overnight on RNAi plates corresponding to knockdowns of individual nucleoporins containing 4.5 mM phenformin. After about 60 hours, about 10 L4 worms were picked onto an agar pad for imaging. Empty vector RNAi plus either phenformin treatment and water served as positive and negative controls, respectively. For the starvation RNAi screen, WBM392 worms were synchronized by egg-laying overnight on RNAi plates corresponding to knockdowns of individual nucleoporins. After about 60 hours, about 20 L4 worms were picked onto unseeded RNAi plates. After a starvation period of 4 hours, about 10 worms were picked onto an agar pad for imaging. Three biological replicates were performed for each RNAi clone.

### Nile Red staining

Neutral lipids were stained and quantified by Nile Red staining as described^98^. Briefly, Nile Red (Fisher Scientific, Cat# N1142) was dissolved in acetone at a stock solution concentration of 5 mg/ml and kept in the dark. Before staining, the working solution was freshly diluted to 30 μg/ml with 40% isopropanol. About 200 young adult worms were synchronized by egg-laying and collected for staining. After 3x washing by PBST (1x PBS plus 0.01% Trion X-100) to remove as much bacteria as possible, the worms were fixed by incubating with 100 μl 40% isopropanol for 3 min. at room temperature. The fixed worms were stained in 400 μl Nile Red working solution by rotating for 2 hours at room temperature, then centrifuged and suspended in 400 μl 1x PBST. The worm pellet was subject to imaging by a Leica DM6 B microscope with THUNDER Imager after rotating in PBST for 30 min.

### Heterochromatin staining

Heterochromatin in worm intestinal cells was stained by 4’,6’-diamidino-2-phenylindole hydrochloride (DAPI; Sigma, Cat#D9542). About 200 worms at day 1 adulthood were collected and washed by M9 to remove bacteria as much as possible, then were fixed with 0.5 ml methanol at −20 °C for 10 min. After 3x centrifugation and washing with M9, the worm pellet was suspended and incubated with 0.5 ml 100 ng/ml DAPI for 30 min with rotation at room temperature. After 3x washing with M9 and collection with centrifugation, the worm pellet was subjected to further imaging by Zeiss LSM 800 with an Airyscan detector.

### Microscopy

Microscopic images were taken on a Leica DM6 B microscope with THUNDER Imager and a Zeiss LSM 800 confocal microscope with an Airyscan detector. For imaging of living worms, about 10-15 worms at the indicated age were mounted on 5% agar pads and anesthetized using 5 mM levamisole (Sigma-Aldrich, Cat#L9756) for each biological replicate.

To image the *acs-2p::GFP* reporter and *sur-5p::NLS::GFP*, images were captured on a Leica DM6 B microscope with Thunder Imager and a 5x objective (for *acs-2p::GFP* reporter) or an x63 objective (for *sur-5p::NLS::GFP*). To image Nile Red stained worms, the worm pellet was dropped onto the slice and subjected to imaging on a Leica DM6 B microscope with Thunder Imager with a 10x objective. For the subcellular imaging of endogenous *gfp::npp-16*, *mKate2::npp-11,* and DAPI staining, images were captured on a Zeiss LSM 800 confocal microscope with an Airyscan detector and 63x/1.4 Oil DIC M27 objective by focusing on the maximal nuclear diameter in DIC images in the anterior intestinal cells, and the original images were then subjected to Airyscan processing before analysis.

For the fluorescence recovery after photobleaching experiment, photo-bleaching and imaging were performed on a Zeiss LSM 800 confocal microscope with an Airyscan detector and 63x/1.4 Oil DIC M27 objective, and the original images were then subjected to Airyscan processing before analysis. The nuclear area was photo-bleached with a 448 nm laser with 100% power and the images were captured every 5 seconds for 150 seconds.

### Image analysis

For *acs-2p::GFP* quantification, in each biological replicate, about 10 worms were laid on the agar pad side by side, and the mean intensity of the entire worm area was quantified by Image J. For Nile Red quantification, the posterior intestinal cells from 10-15 worms were selected and quantified by Image J for each biological replicate. We used ZEISS ZEN Microscopy Software to quantify the fluorescence of endogenous GFP::3xFLAG::AID::NPP-16 in the anterior intestinal cells from 30-40 worms across three biological replicates. The GFP intensity of the nuclear envelope (NE) is defined as the maximum value of a line drawn across the NE. The GFP intensity of nucleoplasm is defined as the mean value of a line drawn randomly in the nucleoplasm. The background is defined as the mean value of a line drawn randomly in the cytosol. To quantify the fluorescence of *sur-5p::NLS::GFP*, the nuclear areas of anterior intestinal cells were selected and quantified by Image J, each biological replicate containing at least 10 worms, and three replicates were performed. For the quantification of FRAP on *vha-6p::GFP* worms, during the photo-bleaching, another area in the cytosol was selected and analyzed parallelly with the bleached nuclear area by ZEISS ZEN Microscopy Software, which is defined as the background. At least three biological replicates were performed. All fluorescent quantifications were normalized to the mean of the control group.

### qRT-PCR

About 1000 worms were synchronized by bleaching and L1 arrest and then collected into 500 μl RNAzol (Sigma-Aldrich, Cat# R4533) after washing three times with M9. The worms were disrupted with a TissueLyser II (QIAGEN) at the frequency of 25 times/sec for 2 minutes in 500 μl RNAzol and 200 μl RNase-free water. After incubating at room temperature for 15 min, the samples were centrifuged at 12000 g for 15 min. The resulting supernatant was added to an equal volume of isopropanol (about 680 ml) and mixed by vortexing. After incubating at room temperature for 15 min, the RNA was pelleted by centrifuge at 12000 g for 10 min. The RNA pellet was air-dried after twice washing with 400 ml 70% ethanol and finally dissolved in 30 μl RNase-free water. The RNA concentration and quality were then determined by NanoDrop One C Microvolume UV-Vis Spectrophotometer (Thermo Fisher Scientific). The cDNA was generated by QuantiTect Reverse Transcription Kit (QIAGEN, Cat# 205314) following the manufacturer’s instructions. In brief, 800 ng RNA was diluted by RNase-free water to 12 μl and incubated with 2 μl gDNA Wipeout buffer at 42 °C for 2 min to remove gDNA contamination.

After adding 2 μl Quantiscript ® Reverse Transcriptase, 2 μl RT Primer Mix and 4 μl Quantiscript RT Buffer, cDNA was generated by incubating at 42 °C for 30 min and 95 °C for 3 min. The cDNA product was 100x diluted by DNase/RNase-free water as the template for qPCR. The qPCR was performed with QuantiTect® SYBR® Green PCR (QIAGEN, Cat# 204145) following the manufacturer’s instructions. The reaction was performed in 10 μl containing 3 μl diluted cDNA template as indicated above on the CFX96 Real-Time System (Bio-Rad). *act-1* mRNA and *act-1* promoter serve as the internal controls for quantifying the corresponding cDNA and gDNA from ChIP, respectively. Actin mRNA was used as the internal control for quantifying the corresponding cDNA in mammalian cells.

### Immunoprecipitation

About 50000 L4 worms were synchronized by bleaching and rotation in M9 buffer overnight leading to synchronous L1 arrest. After plating on bacteria, worms were collected at the L4 stage by washing them free from plates with M9. The bacteria were removed by three M9 washes and worms were collected at each wash by centrifugation. The worm pellet was suspended in a non-denaturing lysis buffer (20 mM Tris-HCl pH 8.0, 137 mM NaCl, 10% glycerol, 1% Triton-100X, 2mM EDTA, supplemented with protease inhibitor cocktail), and sonicated by Misonix Sonicator 3000 (Misonix) with a power output of 30 W and 10 seconds on / 30 seconds off pulse for eight times on ice. After centrifugation at 16000 g at 4°C for 15 min to clear the lysate of insoluble debris, the supernatant protein concentration was measured by BCA kit. Five mg (in about 2 ml) input was used for further immunoprecipitation (IP). The anti-FLAG beads were added into the input and rotated at 4 °C overnight after blocking by 5% BSA at 4°C for 1 hour. For anti-GFP IP, 2 μg GFP antibody was added into the protein supernatant overnight and then incubated with 50 μl pre-washed protein A beads for 1 hour at 4°C. The beads containing the targeted protein were then washed three times with washing buffer (50 mM Tris-HCl pH7.5, 150 mM NaCl, and 6 mM MgCl_2_, supplemented with cocktail protease inhibitor). After removing the supernatant, the immunoprecipitated protein was boiled with 100 μl 1x Laemmli loading buffer at 95 °C for 10 min, then subjected to western blotting. Additional phosphatase inhibitors (0.1mM Na_3_VO_4_, 5mM beta-glycerophosphate, 1μM okadaic acid sodium salt, 10mM NaF, 1mM sodium molybdate) were supplemented into the lysis buffer and washing buffer for western blots specifically intended to detect phosphorylation levels of target proteins.

### Western blotting

About 5000 worms were synchronized by bleaching and L1 arrest, plated, and collected at the L4 larval stage. After washing three times to remove bacteria, the worm pellet was added into 100 μl RIPA buffer (50 mM Tris-HCl pH7.4, 150 mM NaCl, 1% Triton-100X, 1% sodium deoxycholate, 0.1% SDS, 1 mM EDTA, supplemented with protease inhibitor cocktail).

Additional phosphatase inhibitors (0.1mM Na_3_VO_4_, 5mM beta-glycerophosphate, 1μM okadaic acid sodium salt, 10mM NaF, 1mM sodium molybdate) were used for blots against p-Thr, RxxT/S-p, and p-Pol II^Ser5^. The worms were crushed with a tissue grinder about 100 times. The protein supernatant was clarified by centrifugation at 14,000 g at 4°C for 15 minutes to remove insoluble debris, and protein concentration was determined by BCA kit (Thermo Fisher Scientific, Cat# 23225). After adding 4x loading buffer (Bio-Rad, Cat# 1610738) containing 10% β-mercaptoethanol, and boiling at 95 °C for 10 minutes, protein in the supernatant was separated by SDS-PAGE (Bio-Rad, Cat# 456-1084) and transferred electrophoretically to nitrocellulose membranes (Bio-Rad, Cat#1620115). Blots were blocked with 5% non-fat milk or BSA (for detection of phosphorylation) in TBST to reduce background. The membrane was blotted with primary antibody against FLAG (1:3000 dilution, Sigma-Aldrich, Cat# M8823), β-actin(1:5000 dilution, Abcam, Cat# ab14128), GFP (1:3000 dilution; Cell Signaling Technology, Cat# 2956), p-Thr (1:2000 dilution, Cell Signaling Technology, Cat# 9381), p-RXXS/T (1:2000 dilution, Cell Signaling Technology, Cat# 9614), p-AMPK^Thr172^ (1:2000 dilution, Cell Signaling Technology, Cat# 50081), p-Pol II^Ser5^ (1:2000 dilution, BioLegend, Cat# 904001), and RFP (1:2000 dilution, Bulldog-Bio, Cat# RMA6G6) at 4°C overnight, and subsequently the appropriate secondary antibody, either Goat anti-Rabbit IgG (1:5000 dilution, Thermo Fisher Scientific, Cat# 31462) or Goat anti-Mouse IgG (1:5000 dilution, Thermo Fisher Scientific, Cat# G-21040). The blotting signals were captured with the ChemiDoc™ MP Imaging System (Bio-Rad) and quantified by ImageJ.

### RNA sequencing and data analysis

#### RNA extraction

About 1000 worms at day 1 adulthood of each genotype (WT, *npp-16OE,* and *npp-16*^Δ*IDR*^*OE*) were synchronized by egg-laying and collected into 500 μl RNAzol (Sigma-Aldrich, Cat# R4533) after 3x washing for total RNA extraction in biologically independent quadruplicates (12 total samples). The total RNA was extracted, and the genomic DNA was removed using the Direct-zol RNA Miniprep Plus Kit (ZYMO research, Cat# R2072) following the manufacturer’s instructions.

#### RNA sequencing

The extracted total RNA was evaluated for quality control using a NanoDrop One C Microvolume UV-Vis Spectrophotometer (Thermo Fisher Scientific), with an A260/A280 > 1.9 and an A260/A230 > 2.2. Samples were sent to Azenta (Genewiz) for additional quality control, library preparation, and mRNA sequencing. Samples were first validated for RNA integrity with a RIN score > 8 and DV200 > 70 using an RNA Tapestation 4200 (Agilent). Illumina library preparation was performed using polyA selection for mRNA species. Approximately 20 million paired-end 150bp reads were generated per sample, with ≥ 80% of bases passing a Phred quality score ≥ Q30.

#### Data analysis

Fastq read quality control, adapter trimming, quality score filtering, and quasi-alignment were all performed using custom UNIX/Bash shell scripts on the Mass General Brigham ERISTwo Scientific Computing CentOS 7 Linux Cluster. Reads were analyzed for quality control using FastQC v0.11.8 (http://www.bioinformatics.babraham.ac.uk/projects/fastqc) and MultiQC v1.19^99^. Reads were then filtered for Illumina adapter contamination, truncated short reads or low-quality base calls using BBDuk (BBTools)^100^. The subsequently trimmed and cleaned reads were then quasi-aligned to the *Caenorhabditis elegans* reference transcriptome annotation (WBcel235, Ensembl Release 105) using Salmon v1.9.0^101^, correcting for GC content and sequencing bias using the command parameters ‘--gcBias’ and ‘--seqBias’. All statistical analyses and visualizations were generated using the R v4.3.2 (www.r-project.org) Bioconductor v3.18^102^ statistical environment on a local machine through Jupyter Notebook v6.4.10 (https://jupyter.org). Quasi-aligned transcript quantification files for each sample were collapsed into gene-level count matrices using R package tximport v1.30.0^103^, and paired differential expression was calculated using R package DESeq2 v1.42.1^104^ with a design formula of ‘∼ Genotype’. Genes were considered differentially expressed with a Benjamini-Hochberg False Discovery Rate (FDR) corrected P value < 0.05 and an absolute log2 transformed fold change of 1^105^. Volcano plots (Figure 6E) were generated using R package ggplot2 v3.5.0, heatmaps were generated using R package pheatmap v1.0.12, and Venn diagrams (Figure S6A) were generated using R package ggvenn v0.1.10. Gene set overrepresentation analyses for genes identified as differentially expressed (with thresholds set as indicated above) between *npp-16OE* and *npp-16*^Δ*IDR*^*OE* were performed using R package ReactomePA v1.46.0 for Reactome-based pathway terms^75,76^(Figure 6F). Visualizations were post-edited for font, sizing, and appearance using Adobe Illustrator and the Adobe Creative Cloud Suite. All custom bash scripts, Jupyter Notebook files, and processed fastq and transcript count files used in these analyses will be provided upon reasonable request by the corresponding author. All raw fastq files and gene count matrices have been uploaded to NCBI Gene Expression Omnibus (GEO) and can be retrieved through the accession number ‘GSE268357’.

#### Chromatin immunoprecipitation

Chromatin immunoprecipitation (ChIP) was performed as reported with minor modifications^106^. In brief, worms were synchronized by bleaching and L1 arrest, about 50000 arrested L1s were dropped onto the NGM seeded with OP50 in ten 10cm dishes and collected after two days when they grew to the L4 larval stage. After washing three times with M9 to remove as much bacteria as possible, the worms were incubated in crosslinking buffer (PBS containing 1% formaldehyde (vol/vol)) for 20 min at room temperature. The cross-linking was then quenched by adding 200 μl 2.5 M glycine and incubation for another 20 min. After washing three times with PBS containing protease inhibitor cocktail (Sigma-Aldrich, Cat# 11873580001), and additional phosphatase inhibitors were added (0.1mM Na_3_VO_4_, 5mM beta-glycerophosphate, 1μM okadaic acid sodium salt, 10mM NaF, 1mM sodium molybdate) only for p-Pol II^Ser5^ ChIP. The worm pellet was then suspended in 1 ml lysis buffer (50 mM HEPES-KOH, pH 7.5, 150 mM NaCl, 1 mM EDTA, 0.1% (wt/vol) sodium deoxycholate, 1% (vol/vol) Triton X-100, 0.1% (wt/vol) SDS, protease inhibitor cocktail). The lysate was split into 500 μl aliquots in 1.5 ml tubes and then sonicated by Misonix Sonicator 3000 (Misonix) with a power output of 30 W and 10 seconds on / 30 seconds off pulse for eight times on ice. After 16,000 g centrifugation at 4°C for 15 min, protein concentration in the cleared supernatant was measured by BCA kit (Thermo Fisher Scientific, Cat# 23225). After being adjusted to 2 mg/ml with lysis buffer, the supernatant was split into 1 ml for each reaction and 50 μl of it was kept at −80 °C as the input sample. The antibody was added into 1 ml (2 mg protein) supernatant and incubated at 4 °C overnight. 5 μg anti-GFP antibody was used for each reaction for the ChIP in *npp-16OE* and *npp-16^IDR^OE* samples, 10 μg anti-GFP was used for the ChIP in endogenous *GPF::AID::3xFLAG::npp-16* samples due to its low abundance, and 5 μg anti-p-Pol II-Ser5 was used for the ChIP in *npp-16OE* and *npp-16*^Δ*IDR*^*OE* samples. After the antibody incubation, 50 μl slurry of Protein A/G magnetic agarose beads (Thermo Fisher Scientific, Cat# 78610) were added after pre-washing with lysis buffer three times in order to capture the antibody and target protein and incubated for 1 hour at 4 °C. The magnetic beads were washed twice with W1 buffer (50 mM HEPES-KOH, pH 7.5, 150 mM NaCl, 1 mM EDTA pH 8.0, 1% (wt/vol) sodium deoxycholate, 1% (vol/vol) Triton X-100, 0.1% (wt/vol) SDS, and protease inhibitor cocktail), twice with W2 buffer (50 mM HEPES-KOH, pH 7.5, 1 M NaCl, 1 mM EDTA, pH 8.0, 0.1% (wt/vol) sodium deoxycholate, 1% (vol/vol) Triton X-100, 0.1% (wt/vol) SDS, and protease inhibitor cocktail), once with W3 buffer (50 mM Tris-Cl, pH 8.0, 0.25 mM LiCl, 1 mM EDTA, 0.5% (vol/vol) NP-40 and 0.5% (wt/vol) sodium deoxycholate), and three times with 1xTE (10 mM Tris-Cl, pH 8.0, 1 mM EDTA in water). The protein on the magnetic beads was then digested with 40 μg proteinase K (Thermo Fisher Scientific, Cat# 26160) at 45 °C for 2 hours, and the protein in 50 μl input was digested by 50 μg proteinase K at 55 °C for 4 hours. Cross-links were reversed by incubation at 65 °C overnight, releasing the DNA fragments. The DNA in each sample was extracted twice with 250 μl phenol–chloroform–isoamyl alcohol (Sigma-Aldrich, Cat# P3803) and centrifuged at 14,000 g for 10 minutes at room temperature. The DNA was then pelleted in 1 ml ethanol containing 1 μg glycogen (Thermo Fisher Scientific, Cat# R0551) with incubation at −80 °C for 20 minutes and centrifuged at 14,000 g for 10 minutes. After washing with 500 μl 70% ethanol, the DNA pellet was air-dried briefly, and dissolved in 200 and 40 μl elution buffer (10 μM Tris-HCl, pH 8.5) for input and ChIP samples, respectively. For downstream reactions, the DNA was 10x diluted with water and used as the template in subsequent PCR reactions.

## Statistical analysis

The RNA-seq analysis was performed as described above. Other statistical analyses were performed by GraphPad Prism with the indicated methods described in figure legends. Detailed raw quantification and statistics for all data in this manuscript are shown in Tables S1 and S2. Asterisks shown in the figures represent the significance * p<0.05, ** p<0.01; *** p<0.001; **** p<0.0001, ns: non-significant.

## Acknowledgments

We thank Dr. William Mair and Dr. Heng-Chi Lee for the kind gift of worm strains. We thank members of Soukas lab for the helpful discussions and critical reading of the manuscript. This work was supported by NIH/NIA grants R01AG058256 and R01AG069677 (to AAS), the Weissman Family MGH Research Scholar Award (to AAS), and a gift from Stuart and Suzanne Steele and the Obesity Research Fund at the Mass General Research Institute Center for Genomic Medicine. This research was conducted in part while Y.Z. was a Glenn Foundation for Medical Research Postdoctoral Fellow. Thank the NIH/NIDDK-funded NORC of Harvard (P30DK040561) and the NIH/NIDDK-funded Boston-Area DERC (P30DK135043) for core services. Some worm strains in this study were provided by the Caenorhabditis Genetics Center (CGC) funded by the NIH Office of Research Infrastructure Programs (P40OD010440).

## Author contributions

Conceptualization, Y.Z. and A.A.S.; Methodology, Y.Z. and A.A.S.; Formal analysis, Y.Z. and F.A.; Investigation, Y.Z. and F.A.; Resources, Y.Z. and A.A.S.; Writing-Original Draft, Y.Z. and A.A.S; Writing-Review & Editing, Y.Z., F.A. and A.A.S.; Supervision, A.A.S.; Funding Acquisition, A.A.S.

## Competing Interests

Alexander A. Soukas has financial interests in Atman Health, LLC, a company developing an AI-based platform for remote clinical care. The interest of Dr. Soukas was reviewed and is managed by MGH and Mass General Brigham in accordance with their conflict of interest policies.

