## Supplemental Figures for "The nuclear pore complex connects energy sensing to transcriptional plasticity in longevity"

### Supplementary Information

#### Supplementary Information contains seven figures and three tables.

Fig. S1 Zhou et al.

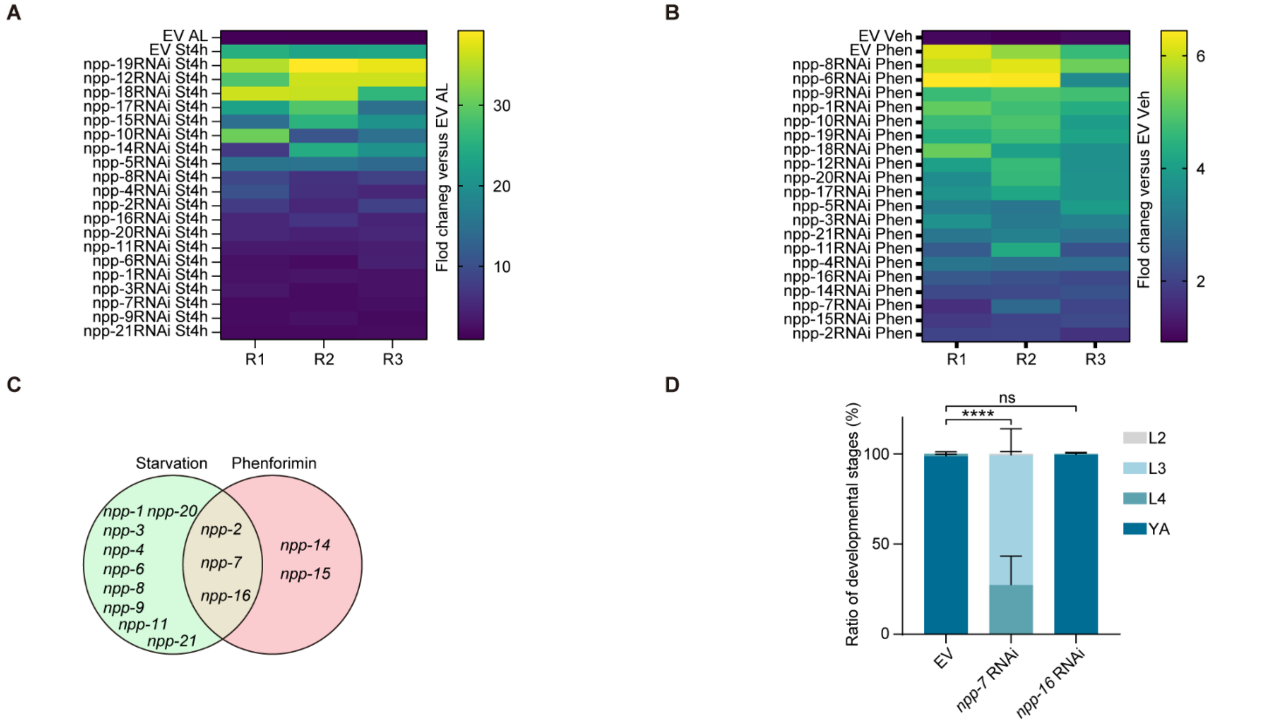


**Figure S1. Detailed results from the nucleoporin RNAi screen on *acs-2p::GFP* reporter , related to Figure 1.**

**A.** Heatmap of *acs-2p::GFP* reporter intensity when treated with starvation for 4 hours (St4h) and the indicated RNAi. *ad libitum* feeding (AL) with empty vector (EV) RNAi serves as a control for normalization. n=3 independent experiments.

**B.** Heatmap of *acs-2p::GFP* reporter intensity treated with phenformin and the indicated RNAi. Treatment with vehicle (Veh) and EV RNAi serves as a control for normalization. n=3 independent experiments.

**C.** Venn diagram showing the overlapping hits from the nucleoporin RNAi screen results in Figures 1A and 1B, with the green and red circles representing the RNAi strains blunting *acs-2p::GFP* induction by starvation and phenformin treatment respectively.

**D.** Percentages of developmental stages in worms treated with indicated RNAi for 56 hours after egg laying. L2: the second larval, L3: the third larval, L4: the fourth larval, YA: young adulthood. n=3 independent experiments.

Bars represent mean ± SD. Statistical significance was determined by two-way ANOVA versus the L4 ratio of worms treated with EV. n=3 independent experiments. ns: non-significant, *p*****<0.0001.

Fig. S2 Zhou et al.

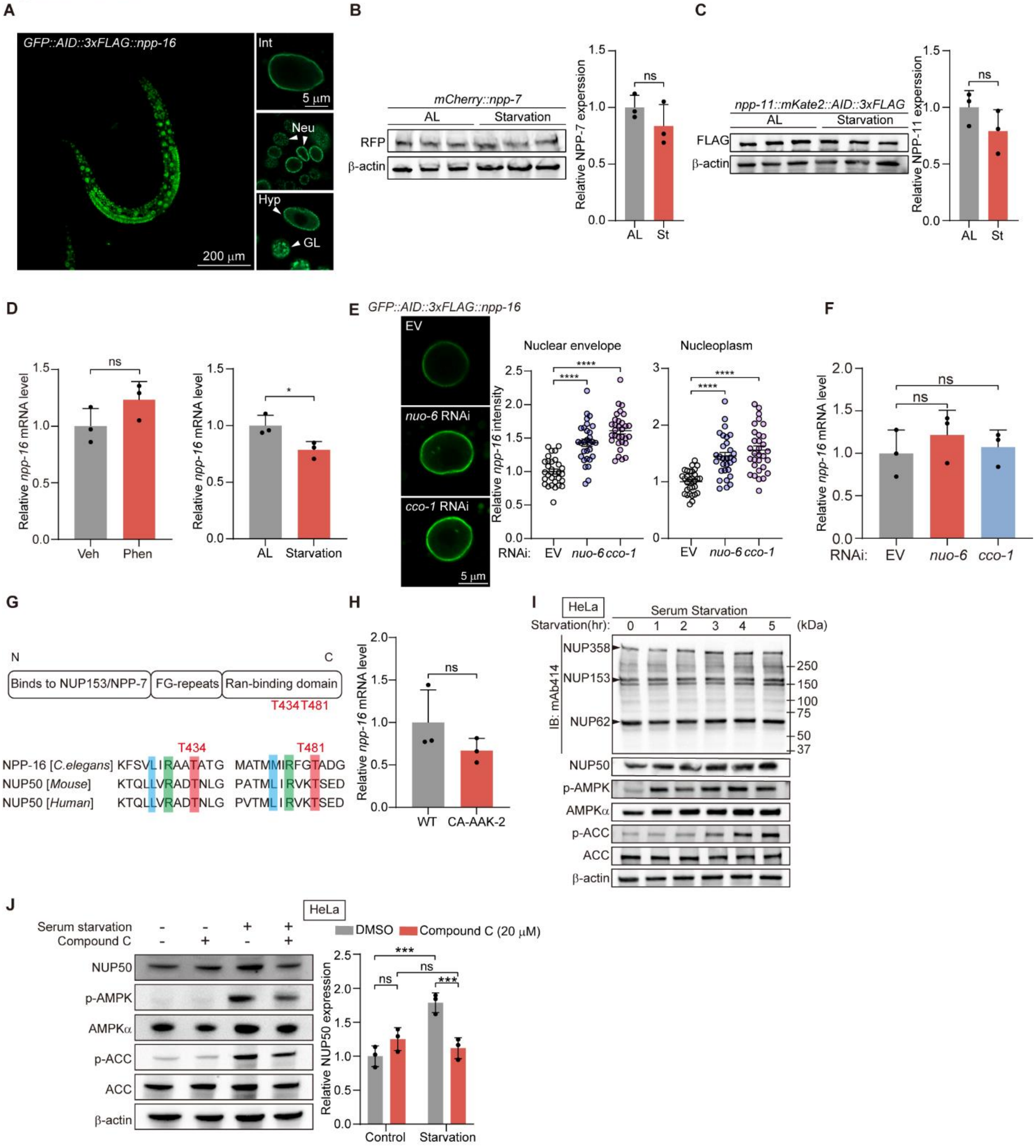


**Figure S2. NPP-16/NUP50 is post-translationally upregulated by nutrient and energetic stress, related to Figure 2.**

**A.** Representative images of endogenous NPP-16 expression pattern in *GFP::AID::3xFLAG::npp-16* worms generated by CRIPSR/Cas9. Int: intestine; Neu: head neuron; Hyp: hypodermis; GL: Germline. Scale bars: 200 μm and 5 μm.
**B.** Endogenous expression of mCherry::NPP-7 is unchanged by starvation for 4 hours. n=3 independent experiments.
**C.** Endogenous expression of NPP-11::mKate2::AID::3xFLAG is unchanged by starvation for 4 hours. n=3 independent experiments.
**D.** qRT-PCR analyses of *npp-16* in WT worms at L4 with indicated treatment. n=3 independent experiments.
**E.** Fluorescence imaging reveals that endogenous *GFP::AID::3xFLAG::npp-16* is increased by ETC inhibition in the nucleoplasm and on the nuclear envelope of anterior intestinal cells at L4. Scale bars: 5 μm. n=3 independent experiments containing at least 30 worms.
**F.** qRT-PCR analyses reveal that *npp-16* mRNA level is unchanged by ETC inhibition at L4. n=3 independent experiments.
**G.** The consensus motif of AMPK phosphorylation is conserved between human NUP50, mouse NUP50, and nematode NPP-16.
**H.** *npp-16* mRNA levels are unchanged by CA-AAK-2 by qRT-PCR. n=3 independent experiments.
**I.** Human NUP50 is induced by serum starvation in HeLa cells at the indicated times, whereas NUP153 and NUP62 are not. mAb414 is an antibody recognizing multiple FG-domain-containing nucleoporins. The phosphorylation of $ACC^{Ser79}$ (p-ACC) and $AMPK^{Thr172}$ (p-AMPK) serves as positive controls for energetic stress activation.
**J.** Treatment with 20 μM of the AMPK inhibitor Compound C prevents the induction of human NUP50 by serum starvation for 3 hours in HeLa cells. DMEM supplemented with 10% FBS serves as the negative control (Ctrl) for serum starvation. n=3 independent experiments.
Bars represent mean ± SD (B-D, F, H, and J) and mean ± SEM (E). Statistical significance was determined by unpaired *t*-test (B-D, and H), One-way ANOVA (E and F), and two-way ANOVA (J). Relative protein and mRNA levels were normalized to β-actin (B, C, and J) and *act-1* (D, F, and H) respectively. Empty vector (EV) serves as the negative control for RNAi experiments. ns: non-significant, *$p$<0.05, ***$p$<0.001, ****$p$<0.0001.

Fig. S3 Zhou et al.

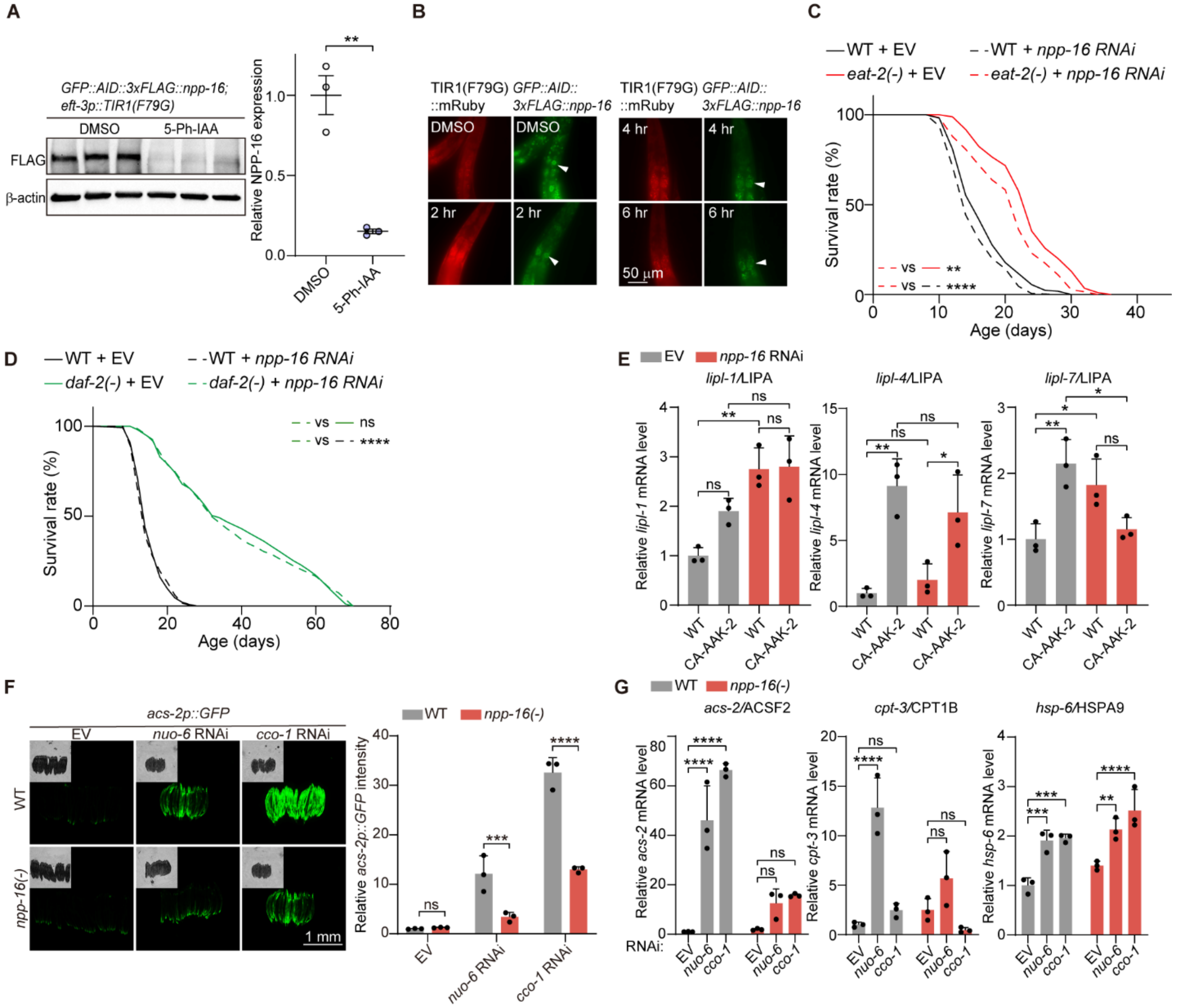


**Figure S3. NPP-16/NUP50 is necessary for pro-longevity paradigms associated with energetic stress, related to Figure 3.**

**A.** Endogenous GFP::AID::3xFLAG ::NPP-16 is degraded efficiently by the AID-TIR1(F79G) system as confirmed by immunoblotting. Worms were treated with 5 μM 5-Ph-IAA from hatching to L4. n=3 independent experiments.

**B.** Representative images of *gfp::AID::3xFLAG::npp-16; eft-3p::AtTIR1(F79G)::mRuby* treated with 5 μM 5-Ph-IAA for the indicated times at day 1 of adulthood (D1). The intestinal nucleus is indicated by white arrowheads. Scale bar: 50 μm.

**C-D.** Lifespan analyses reveal that the lifespan extension of *eat-2* and *daf-2* mutants is unaffected by *npp-16* RNAi.

**E.** qRT-PCR analyses reveal that increased *lipl-1*, *lipl-4,* and *lipl-7* mRNA expression by CA-AAK-2 is unchanged by *npp-16* RNAi. n=3 independent experiments.

**F.** Fluorescence imaging reveals that *npp-16(-)* mutation suppresses the induction of *acs-2p::GFP* when the ETC is inhibited by *nuo-6* and *cco-1* RNAi. Scale bar: 1 mm. n=3 independent experiments.

**G.** qRT-PCR analyses reveal that the increased expression of *acs-2* and *cpt-3* by ETC inhibition is rescued in *npp-16(-)* mutants, however, *npp-16(-)* mutation has no effect on the induction of *hsp-6* by ETC inhibition. n=3 independent experiments.

Bars represent mean ±SD. Statistical significance was determined by unpaired *t*-test (A), one-way ANOVA (E), or two-way ANOVA (F and G). Empty vector (EV) serves as the negative control for RNAi experiments. Relative protein and mRNA levels were normalized to β-actin (A) and *act-1* (E and G) respectively. ns: non-significant, *$p<0.05$, **$p<0.01$, ***$p<0.001$, ****$p<0.0001$. See also Table S1 for independent biological replicates and summary lifespan statistics.

Fig. S4 Zhou et al.

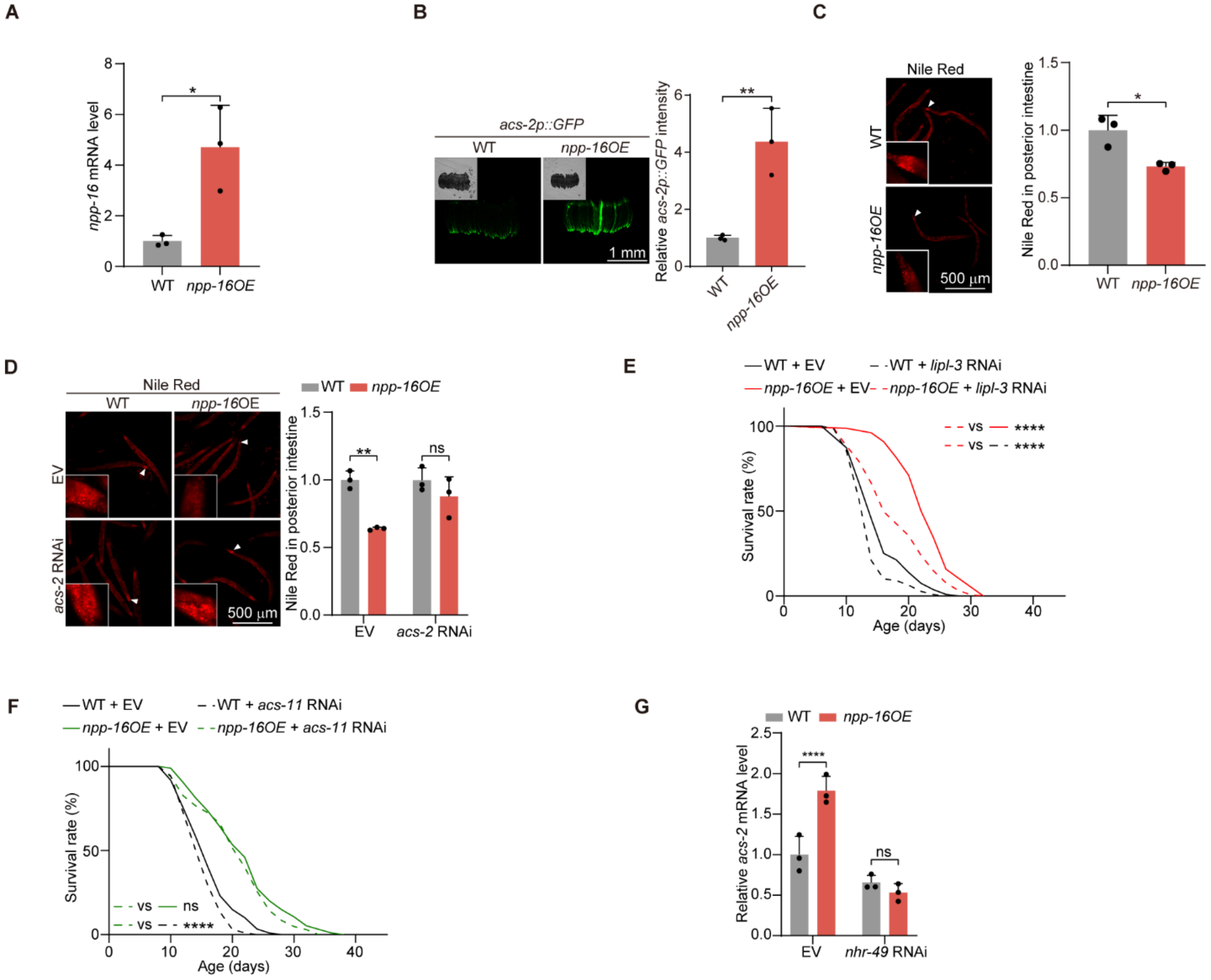


**Figure S4. NPP-16/NUP50 promotes longevity by activating lipid catabolism, related to Figure 4.**

**A.** qRT-PCR validation of the level of overexpression of *npp-16* mRNA in *npp-16OE* transgenics. n=3 independent experiments.

**B.** Fluorescence imaging reveals that *npp-16OE* increases expression of the *acs-2p::GFP* reporter at day 1 of adulthood. Scale bar: 1 mm. n=3 independent experiments.

**C.** Neutral lipid storage indicated by fixative Nile red staining reveals that *npp-16OE* decreases the fat mass of posterior intestinal cells, which are indicated by white arrowheads and enlarged in white boxes. Scale bar: 500 μm. n=3 independent experiments containing at least 60 worms per group.

**D.** Neutral lipid storage indicated by fixative Nile red staining reveals that *acs-2* RNAi rescues the fat mass decrease in *npp-16OE* worms. Intestinal cells are indicated by white arrowheads and enlarged in white boxes. Scale bar: 500 μm. n=3 independent experiments containing at least 45 worms per group.

**E**. Knockdown of *lipl-3* by RNAi partially suppresses lifespan extension seen in *npp-16OE* transgenic worms.

**F.** Knockdown of *acs-11* by RNAi does not affect the longevity phenotype of *npp-16OE* worms.

**G.** qRT-PCR validation shows that *nhr-49* RNAi blunts increase in *acs-2* mRNA in *npp-16OE* worms. n=3 independent experiments.

Bars represent mean ± SD. Statistical significance was determined by unpaired *t*-test (A-C) and Two-way ANOVA (D and G). Empty vector (EV) serves as the negative control for RNAi experiments. See also Table S1 for independent biological replicates and summary lifespan statistics. Relative mRNA levels were normalized to *act-1* (A and G). ns: non-significant, *$p<0.05$, **$p<0.01$, ****$p<0.0001$.

Fig. S5 Zhou et al.

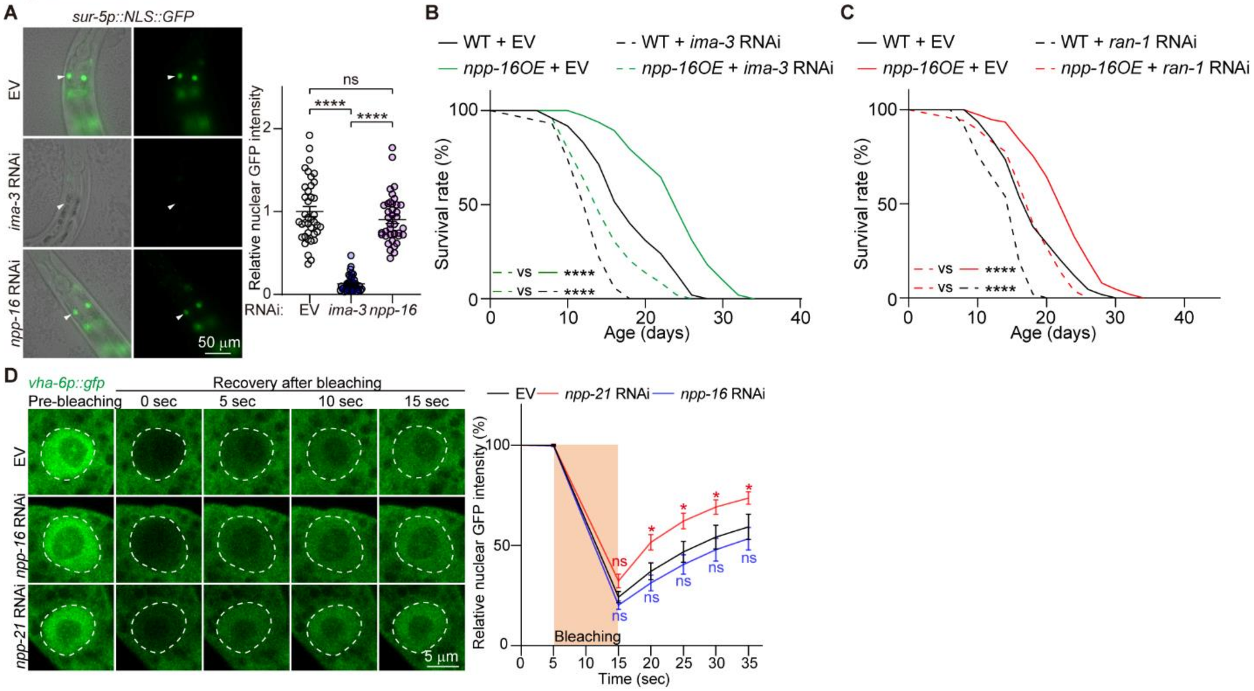


**Figure S5. NPP-16/NUP50 does not impact nuclear transport and permeability in *C. elegans*, related to Figure 5.**

**A.** The nuclear GFP intensity of *sur-5p::NLS::GFP* in the anterior intestinal cells of day 1 adult worms treated with the indicated RNAi reveals that *npp-16* RNAi does not alter active nuclear transport. *ima-3* RNAi serves as a positive control. Scale bar: 50 µm. n=3 independent experiments containing at least 39 worms per group.

**B.** *npp-16OE* extends the lifespan of worms treated with *ima-3* RNAi from hatching.

**C.** *npp-16OE* extends the lifespan of worms treated with *ran-1* RNAi post-developmentally.

**D.** Knockdown of *npp-16* by RNAi does not change the fluorescence recovery rate of intestinal GFP after photo-bleaching, a proxy for passive nuclear transport. Dashed circles indicate the nuclear areas. *npp-21* RNAi serves as a positive control. Scale bar: 5 µm. n=3 independent experiments.

Bars represent mean ± SEM (A) and mean ± SD (D). Statistical significance was determined by one-way ANOVA (A) and two-way ANOVA (D). Empty vector (EV) serves as the negative control for RNAi experiments. See also Table S1 for independent biological replicates and summary lifespan statistics. ns: non-significant, *$p<0.05$, ****$p<0.0001$.

**Fig. S6 Zhou et al.**

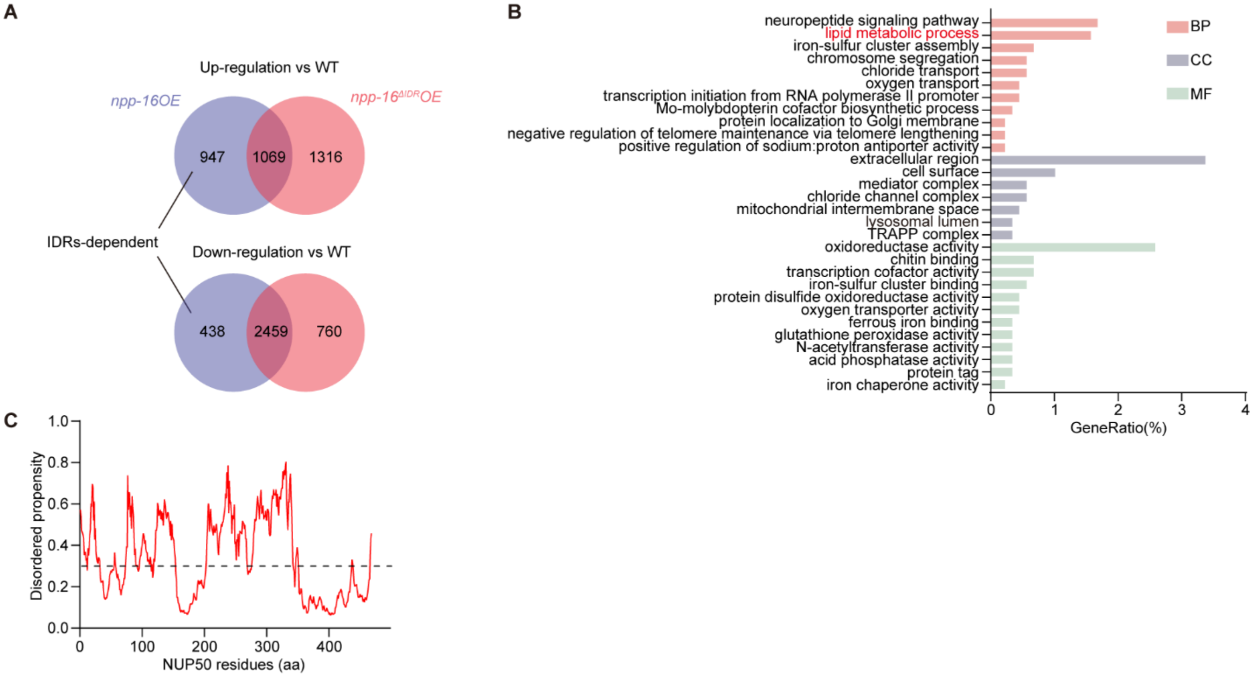


**Figure S6. Transcriptomic alterations of *npp-16OE* and *npp-16*$^{\Delta IDR}$*OE*, related to Figure 6.**

**A.** Venn diagram showing the number of DEGs in *npp-16OE* and *npp-16*$^{\Delta IDR}$*OE* worms versus WT worms. The DEGs are defined by an adjusted *P* value threshold of 0.05 and an absolute $\log_2$ fold-change >1. See also Table S3 for detailed statistics.

**B.** GO-term overrepresentation analysis of the 947 IDR-dependent up-regulated DEGs in *npp-16OE* worms. BP: biological processes; CC: cellular components; MF: molecular functions. Lipid metabolism pathway overrepresentation is highlighted in red. See also Table S3 for detailed statistics.

**C.** The bioinformatic prediction of the intrinsically disordered region (IDR) in human NUP50 by flDPnn, the threshold of 0.3 predictive of an intrinsically disordered region is indicated by a black dashed line.

**Fig. S7 Zhou et al.**

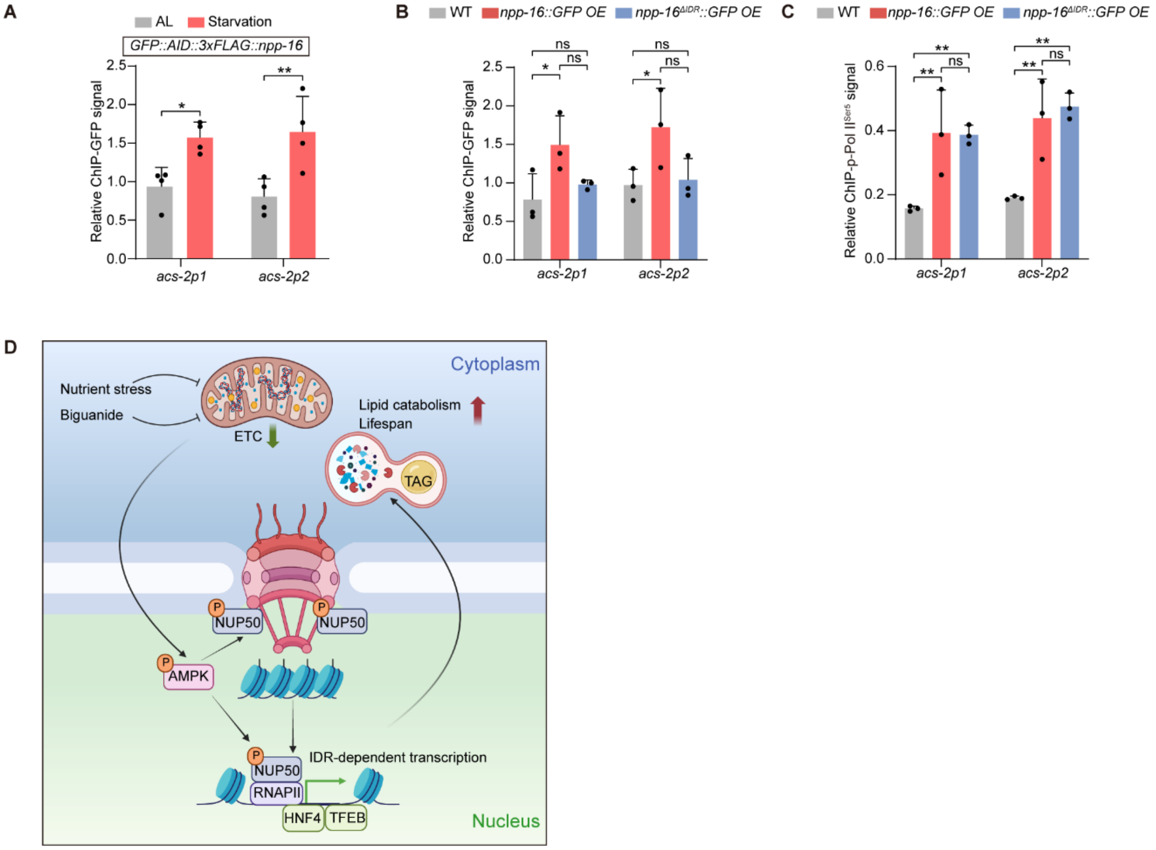


**Figure S7. NPP-16/NUP50 binds to the *acs-2* promoter in an IDR-dependent manner, related to Figure 7.**

**A.** GFP ChIP-qPCR of two regions at *acs-2* promoter (*acs-2p1* and *acs-2p2*) in *GFP::3xFLAG::AID::npp-16* worms treated with *ad libitum* feeding (AL) or starvation for 4 hours at L4. n=4 independent experiments.

**B.** GFP::NPP-16 in *npp-16OE* worms binds to the *acs-2* promoter, whereas GFP::NPP-16$^{\Delta IDR}$ does not. n=3 independent experiments.

**C.** Both *npp-16OE and npp-16*$^{\Delta IDR}$*OE* promote the p-Pol II$^{Ser5}$ recruitment onto the *acs-2* promoter. n=3 independent experiments.

**D.** Schematic depiction of NPP-16/NUP50 bridging energy sensing and adaptive transcription of lipid catabolic genes to modulate organismal metabolism and pro-longevity outcomes. ETC: electron transport chain, TAG: triacylglycerol. Created in BioRender. Zhou, Y. (2023) BioRender.com/v74r321.

Bars represent mean ± SD. Statistical significance was determined by two-way ANOVA. Relative ChIP signals were normalized to the *act-1* promoter (A-C). ns: non-significant, *$p<0.05$, **$p<0.01$.

**Table S1. Lifespan statistics and biological replicates in this study, related to Figure 3, 4, 5, 7, S3, S4, S5, and STAR Methods.**

**Table S2. Detailed statistics of all non-lifespan related assays in this study, related to STAR Methods.**

**Table S3. RNA sequencing analyses of WT, *npp-16OE*, and *npp-16*$^{\Delta IDR}$*OE* worms, related to Figures 6 and S6.**
